# Disentangling the impacts of abiotic and biotic environmental factors and dispersal dynamics on the pangenome fluidity of bacterial pathogens

**DOI:** 10.1101/2025.03.27.645760

**Authors:** Ying-Xian Goh, FNU Hardeep, Hailong Zhang, Jingqiu Liao

**Affiliations:** Department of Civil and Environmental Engineering, Virginia Tech, Blacksburg, VA 24061, United States; Center for Emerging, Zoonotic, and Arthropod-Borne Pathogens, Virginia Tech, Blacksburg, VA 24061, United States; Global Change Center, Virginia Tech, Blacksburg, VA 24061, United States; Department of Computer Science, Virginia Tech, Blacksburg, VA 24061, United States; Department of Business Information Technology, Virginia Tech, Blacksburg, VA 24061, United States

## Abstract

Understanding how pangenomes originate and evolve is crucial for predicting evolutionary trajectories and uncovering ecological interactions of bacterial pathogens. Pangenome fluidity has been attributed to adaptive evolution, yet the underlying ecological drivers for bacterial pathogens persisting in natural reservoirs remain poorly understood. *Listeria monocytogenes* (*Lm*), a foodborne pathogen causing fatal listeriosis, serves as an ideal model for investigating the ecological mechanisms underlying pangenome fluidity in bacterial pathogens due to its high evolutionary divergence, broad ecological versatility, and significant public health concern. Through pangenome analysis of 177 *Lm* isolates representing three evolutionary lineages (I, II, and III) that we isolated from soils across the United States, we found that substantial genome variation was strongly associated with climatic factors (e.g. precipitation and temperature), soil properties (e.g. aluminum, pH, and molybdenum), and bacterial community composition, particularly Nitrospirae, Planctomycetes, Acidobacteria, and Cyanobacteria. These factors exerted selective pressure across many gene functions, with pronounced effects on genes involved in cell envelope synthesis, defense mechanisms, and replication, recombination, and repair. Among *Lm* lineages occupying varied habitats, distinct pangenome properties were observed. Lineage III exhibited a highly fluid pangenome, which was attributed to local adaptation to nutrient-limited conditions and strong dispersal limitation. In contrast, lineage I maintained a conserved pangenome, likely due to frequent homogenizing dispersal. Consistent with these dispersal patterns, we identified an elevated risk of soil-to-human transmission in lineage I, evidenced by epidemiological links between three soil-derived and 17 clinical isolates. Collectively, this study reveals the pivotal role of environmental selection imposed by both abiotic factors and bacterial communities in governing the adaptive pangenome evolution in bacterial pathogens. It also highlights significant differences in pangenome flexibility, ecological niches, and transmission dynamics across lineages of the same pathogen species, underscoring the need for tailored source tracking strategies.

**AUTHOR SUMMARY:** Studying the full set of genes found in different strains of a bacterium (i.e. pangenome) helps us understand how bacterial pathogens develop and adapt to changes in the environment. Here, we focused on *Listeria monocytogenes* (*Lm*), a pathogen capable of spreading through food and surviving in diverse environments, to understand how environmental factors and the way that bacteria move across locations can influence the pangenome content in this important bacterium. By analyzing the genomes of 177 *Lm* strains representing three evolutionary lineages (I, II, and III) collected from soils across the United States, we found that variation in climate, soil chemistry, and surrounding bacteria (e.g., Nitrospirae) was closely linked to genetic differences among strains. These environmental conditions seemed to affect genes that help build the cell envelop, protect the bacteria from harm, and fix damaged DNA. We also observed different levels of genome flexibility across *Lm* lineages which were found to be related to how they move across different locations. Lineage III showed evidence of barriers to spreading, which may enhance genetic differentiation across populations, leading to a more flexible pangenome. In contrast, lineage I appeared to spread more readily and was epidemiologically linked to human clinical cases, which may facilitate genetic exchange that reduce pangenome diversity. This study shows that both non-living environmental conditions—like precipitation and pH—and nearby groups of bacteria play a big role in shaping how bacterial pathogens change their genes to survive. It also highlights that different subtypes of the same pathogen can have different gene flexibility and spread in different ways, calling for specific biocontrol measures.

## INTRODUCTION

Pangenome consists of core genes (i.e., genes present in all individuals) and accessory genes (i.e., genes not shared by all individuals), which arise from gene gain and loss through evolutionary processes such as horizontal gene transfer and pseudogene formation [1–3]. Bacterial pathogens vary in pangenome fluidity. Some species, like *Escherichia coli*, have an open pangenome, where the number of genes increases substantially with each newly sequenced genome, while others, such as *Bacillus anthracis*, exhibit a closed pangenome, where the gene count remains stable despite the addition of more genomes [4–6]. The evolutionary mechanisms underlying pangenome fluidity of bacterial pathogens remain a subject of ongoing debate in the field. While some argue that pangenome variation results from neutral evolution in which genetic drift randomly changes the frequency of alleles, our previous study and others suggest it is primarily caused by adaptation [6–8]. Adaptive evolution of pangenomes can be triggered by various environmental pressures that pathogens need to cope with, including abiotic factors, such as antibiotic exposure, mercury, and aluminum toxicity [9–11], and biotic factors such as bacterial mutualism and competition [11,12], and phage predation [13]. For example, antibiotic exposure were found to drive the acquisition of chloramphenicol and streptomycin resistance genes in *Vibrio cholerae* [9]. Also, mutualism between *Salmonella enterica* and *Methylorubrum extorquens* in co-culture has been shown to reduce the impact of loss-of-function mutations and selectively maintain nitrogen uptake genes in *S. enterica* compared to its monoculture [12]. In addition to environmental selection, pangenome fluidity can also be influenced by dispersal. While dispersal can expose pathogens to new environments and promote the acquisition of novel genes [14,15], it can also constrain pangenome diversity through homogenizing dispersal, where the continuous movement of similar genetic material across populations reduces differentiation that would otherwise emerge from localized environmental pressures [16–18]. Understanding how environmental factors and dispersal influence pangenome properties can shed light on the mechanisms and predictability of the evolution of bacterial pathogens under environmental changes. It can also provide insights into transmission risks across environmental contexts and potential spillover to humans. However, due to a lack of in-depth studies integrating genomics with paired abiotic and biotic environmental data for bacterial pathogens in natural reservoirs, the ecological drivers of the pangenome fluidity of bacterial pathogens persisting in their natural reservoirs remain poorly understood.

*Listeria monocytogenes* (*Lm*) is characterized by high genomic diversification, ecological versatility, and public health relevance, which make it an ideal model for studying the ecological mechanisms governing the pangenome evolution of bacterial pathogens. *Lm* is a Gram-positive, motile, non-spore-forming facultative bacterium commonly found as a saprophyte in soils [19,20]. *Lm* is also a foodborne pathogen that causes rare but fatal listeriosis affecting vulnerable populations (i.e., pregnant women, the elderly, infants, and immunocompromised individuals), with a fatality rate of 20-30%, hospitalization rate of 94%, and an annual illness cost of USD 2.6 billion [21,22]. *Lm* can thrive and transmit across diverse habitats (e.g., soil, water, animals, produce, food processing facilities, and hospital settings) [8,19,22–24]. It has formed four distinct evolutionary lineages (designated as I, II, III, and IV) [25] characterized by substantial intraspecific gene content variation [26–28] and unique ecological roles [23,29]. *Lm* lineage I, II, and III are commonly detected in human clinical cases, food-related sources, and soil environments, respectively, while lineage IV is rare and has been detected in ruminants [8,23,25]. It is known that *Lm* genomes undergo adaptive evolution [8,30–33]. For example, *Lm* has developed stress survival islets (SSI) 1 and 2 to cope with environmental stress, with SSI-1 responsible for tolerating high salt and bile salt concentrations and low pH [30], and SSI-2 for tolerance of alkaline and oxidative stress conditions [31]. Also, in food processing environments where benzalkonium chloride is commonly used as a surface disinfectant, *Lm* strains have been found to acquire tolerance genes to withstand its effects [32,33]. These individual studies, however, only provide a snapshot of how environmental factors influence particular genes in *Lm*, with a focus on food-associated environments. The systematic effects of abiotic and biotic factors and dispersal across environments shaping the pangenome fluidity of *Lm* inhabiting its natural reservoir, the soil, remain largely unknown.

Here, we conducted a comprehensive pangenome analysis of 177 *Lm* isolates obtained from 1,004 soil samples that we previously systematically collected from natural environments across the United States (US) [8]. These isolates represent three lineages, I, II, and III. We examined the relationships between the pangenome variation of *Lm* and 34 abiotic variables capturing geolocation, soil properties, climate, and surrounding land use, as well as bacterial community composition characterized by 16S rRNA gene amplicon sequencing. Using a suite of statistical analyses, including machine learning, we identified strong relationships between climate, soil properties, and bacterial community composition, particularly Nitrospirae, Planctomycetes, Acidobacteria, and Cyanobacteria, and pangenome variation at multiple functional categories. Functional enrichment analysis revealed that abiotic and biotic pressures jointly target genes involved in cell envelope synthesis and regulation, with abiotic and biotic factors additionally acting on genes related to defense mechanisms and replication, recombination, and repair, respectively. We also identified distinct pangenome properties across *Lm* lineages, which were detected in distinct environmental conditions. The pangenome of lineage III is highly fluid, but its genome size appears to be streamlined, which we attribute to local adaptation to nutrient-limited conditions and strong dispersal limitation. In contrast, the pangenome of lineage I is conserved, which is likely contributed by frequent homogenizing dispersal across the US. Informed by this observation, we identified a high risk of transmission from soil to humans in lineage I through comparative genomic analysis of soil and clinical isolates. Overall, this study provides insights into how selective pressures triggered by abiotic factors, bacterial communities, and dispersal dynamics can contribute to the pangenome fluidity of bacterial pathogens.

## RESULTS

### Associations between abiotic environmental factors and *Lm* pangenome

For the 177 *Lm* isolates included in this study, a total of 5,873 orthologous genes were identified, with 2,324 and 3,549 classified as core and accessory genes, respectively (**Supplemental Fig. S1**). The gene richness per genome ranged from 2627 to 3082 (**Supplemental Fig. S1**). To identify key abiotic drivers of the pangenome composition of *Lm*, we performed a series of statistical analyses assessing the relationships between 34 abiotic environmental variables for geolocation, soil properties, climate, and surrounding land use, and gene richness as well as gene presence/absence patterns. Variation partitioning analysis (VPA), combined with a permutation test (see Methods), revealed that abiotic factors altogether significantly explained 14.24% of the gene richness variation (one-sided permutation *P* = 0.01; **Fig. 1A**). Among them, soil property and climatic variables were found to be most influential, accumulatively explaining 10.17% and 9.05% of the variation with other variables, respectively, followed by geolocation (8.79%) and surrounding land use (6.36%; **Fig. 1A**). Of note, the interplay of soil properties and climate explained 5.16% of the variation (**Fig. 1A**). To further assess the relationships between abiotic factors and gene richness at the functional level, VPA stratified by Clusters of Orthologous Groups (COGs) was conducted. Results showed that among the 18 COGs, abiotic factors significantly explained the variation of gene richness for 17 COGs (adjusted one-sided permutation *P* < 0.05 for all), with G (carbohydrate transport and metabolism) and K (transcription) exhibiting over 50% of the variation explained (**Fig. 1B**).

**Figure 1.**
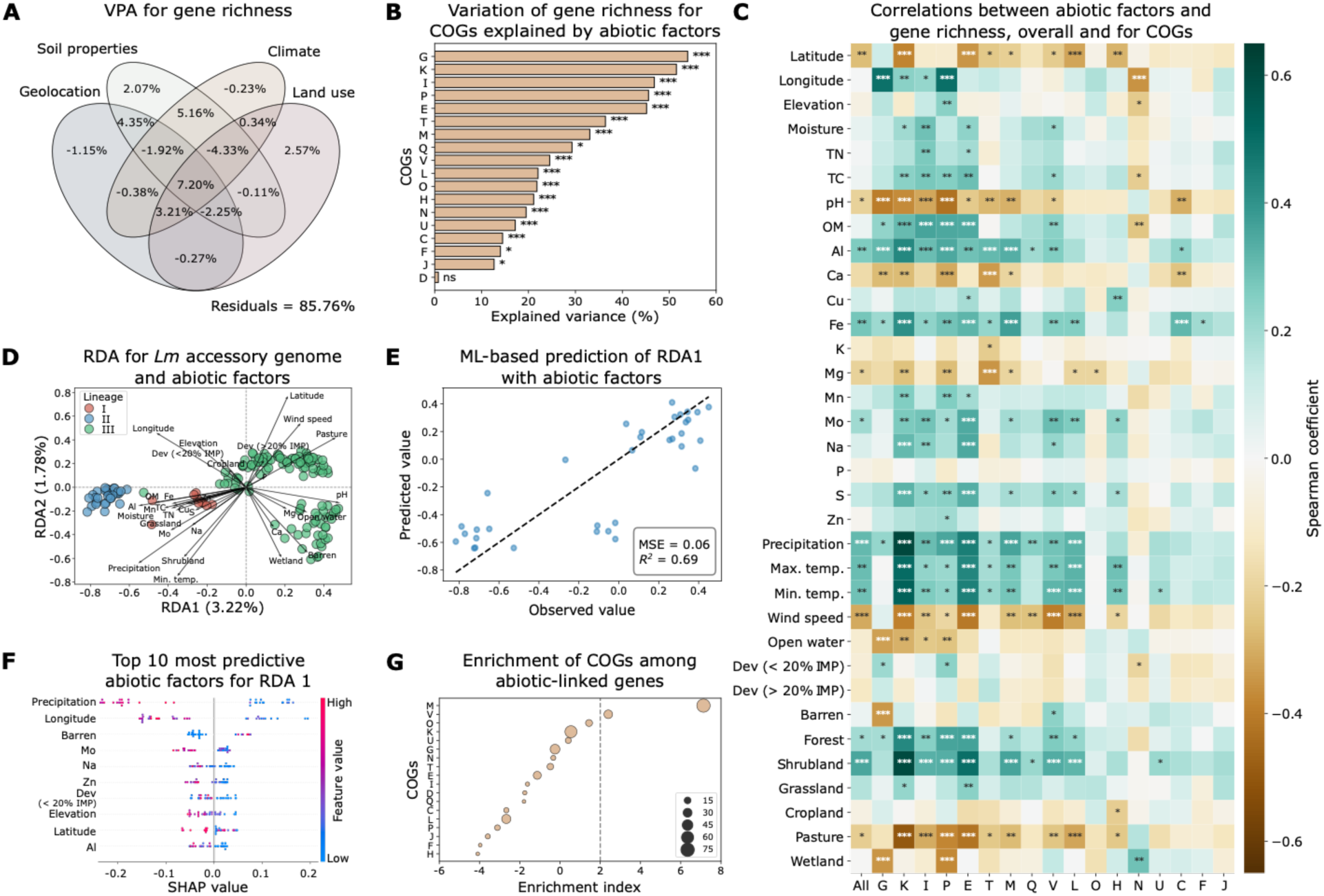
Pangenome fluidity of *Lm* is associated with abiotic factors. **(A)** Venn diagram of variation partitioning analysis (VPA) showing the variation of gene richness explained by geolocation, soil properties, climate, and surrounding land use. Residuals represent unexplained variation. A negative explained variation may be caused by collinearity between predictors and overparameterization of the model. **(B)** Variation of gene richness for COGs explained by abiotic factors in VPA. Statistical significance was assessed using a one-sided permutation test. **(C)** Spearman’s correlation between abiotic factors and gene richness, overall and for significant COGs identified in VPA. Positive and negative correlations are indicated by green and brown, respectively. **(D)** Redundancy analysis (RDA) biplot illustrating the relationships between the *Lm* accessory genome and 31 abiotic variables selected by Lasso regularization. Points represent individual genomes, color-coded by lineages. The length and direction of the arrows indicate the relative importance and correlation of each abiotic variable with the ordination axes. The first (RDA1) and second (RDA2) ordination axes explain 3.22% and 1.78% of the variation of the *Lm* accessory genome explained by selected abiotic factors, respectively. **(E)** Prediction of RDA1 axis values with abiotic factors using the LightGBM machine learning (ML) model. MSE, mean squared error; *R^2^*, coefficient of determination. The dashed line represents the line of perfect agreement (y = x) where predicted values would exactly match observed values. **(F)** The top ten most predictive abiotic factors for RDA1 (SHAP-based; X axis), sorted by descending importance. SHAP values indicate the impact of features on ML model output. **(G)** Enrichment of COGs among abiotic-linked genes. An enrichment index greater than two (grey dashed line) indicates significant overrepresentation (*P* < 0.05). Circle size is proportional to the number of genes annotated to each COG. For **(B)** and **(C)**, significance levels are denoted by “*”, “**”, “***”, and “ns” for adjusted *P* < 0.05, < 0.01, < 0.001, and > 0.05 (not significant), respectively. Abbreviations of COGs for **(B)** and **(G),** and abiotic factors for **(C)**, **(D)**, and **(F)** are described in Methods.

To assess the relationships between individual abiotic factors and gene richness, Spearman’s correlation analysis was performed for overall gene richness as well as that for each COG. For overall gene richness, precipitation, temperature, soil aluminum, iron, and surrounding shrubland coverage were significantly positively correlated with gene richness, while wind speed, magnesium, pH, and surrounding pasture coverage showed a significant negative correlation (adjusted Spearman’s *P* < 0.05 for all; **Fig. 1C**). At the functional level, these abiotic variables were also significantly correlated with gene richness for over half of the 17 significant COGs identified in VPA (adjusted *P* < 0.05 for all; **Fig. 1C; Supplemental Fig. S2**), suggesting a universal effect on the gene content of *Lm* across multiple functions. In contrast, some abiotic variables tend to have a function-specific effect on gene richness. For example, potassium and zinc were only significantly correlated with gene richness for COGs T (signal transduction mechanisms) and P (inorganic ion transport and metabolism), respectively (adjusted *P* < 0.05 for both; **Fig. 1C**).

In addition to gene richness, we assessed the relationships between abiotic variables and the presence/absence patterns of accessory genes. Redundancy analysis (RDA) of 31 out of 34 abiotic variables selected by Lasso regularization showed that 3.22% and 1.78% of the variation in the first (RDA1) and second (RDA2) ordination axis of the accessory gene content, respectively, was explained by these selected variables, with precipitation, minimum temperature, wind speed, aluminum, pH, molybdenum, latitude, longitude, and surrounding shrubland, pasture, barren, and wetland coverage exerting a strong effect (**Fig. 1D**). As the relationships between abiotic variables and accessory gene variance may not be captured by the linear constraints of RDA, we further employed machine learning to model their relationships (see Methods). Results show that with LightGBM and random forests, 69% and 84% of the variation in RDA1 and RDA2 was explained by these selected abiotic variables, respectively (**Fig. 1E; Supplemental Fig. S3A**). Based on Shapley Additive exPlanations (SHAP) analysis [34], precipitation, minimum temperature, aluminum, molybdenum, latitude, longitude, and surrounding coverage of shrubland and barren, which were found to have a strong effect in the RDA biplot (**Fig. 1D**), were also among the top 10 most predictive abiotic factors for RDA1 and/or RDA2 (**Fig. 1F; Supplemental Fig. S3B**). Analyses on gene richness and accessory genomes altogether suggest that the pangenome fluidity of *Lm* is strongly shaped by climatic variables (e.g., precipitation and temperature), soil properties (e.g., aluminum, pH, and molybdenum), and geolocation.

To assess the associations between abiotic factors and individual accessory genes, we compared the difference of abiotic factors for *Lm* genomes with and without a given accessory gene. A total of 803 accessory genes were found to be significantly associated with at least one abiotic factor, which we defined as “abiotic-linked genes” (adjusted Mann-Whitney *U P* < 0.05 for all; **Supplemental Table S1**). Functional enrichment analysis (see Methods) showed that COGs M (cell wall/membrane/envelope biogenesis) and V (defense mechanisms) were significantly enriched among these abiotic-linked genes (**Fig. 1G**). These results along with gene richness analysis stratified by COGs suggest that abiotic factors play a critical role in structuring the gene content of *Lm* across multiple functions, with pronounced effects acting on genes involved in cell envelope formation and defense mechanisms.

### Associations between bacterial community composition and *Lm* pangenome

*Lm* does not exist in isolation within the soil environment; instead, it coexists within communities alongside other bacteria. Among the soil samples positive for *Lm*, a total of 28 phyla were identified. To test the contribution of bacterial community composition (also referred to as “biotic factors”) to the variation of *Lm* pangenome, VPA was performed on the gene richness and the relative abundance of bacterial phyla along with abiotic factors. Bacterial phylum composition alone and cumulatively with abiotic factors significantly contributed to 13.25% and 19.87% of the variation of gene richness, respectively (one-sided permutation *P* < 0.001; **Fig. 2A**). Stratifying by COGs, the variation of gene richness for 14 COGs was significantly explained by both abiotic factors and bacterial phylum composition (adjusted one-sided permutation *P* < 0.05 for all; **Fig. 2B**). Among these COGs, G (carbohydrate transport and metabolism), I (lipid transport and metabolism), P (inorganic ion transport and metabolism), and K (transcription) had over 50% of their variation explained, indicating a significant impact of abiotic and biotic factors on these gene functions (**Fig. 2B**). Of note, COGs L (replication, recombination, and repair), J (translation, ribosomal structure, and biogenesis), and V (defense mechanisms) showed greater variation of gene richness explained by biotic factors than abiotic factors, indicating higher importance of bacterial community composition impacting these functions (**Fig. 2A**).

**Figure 2.**
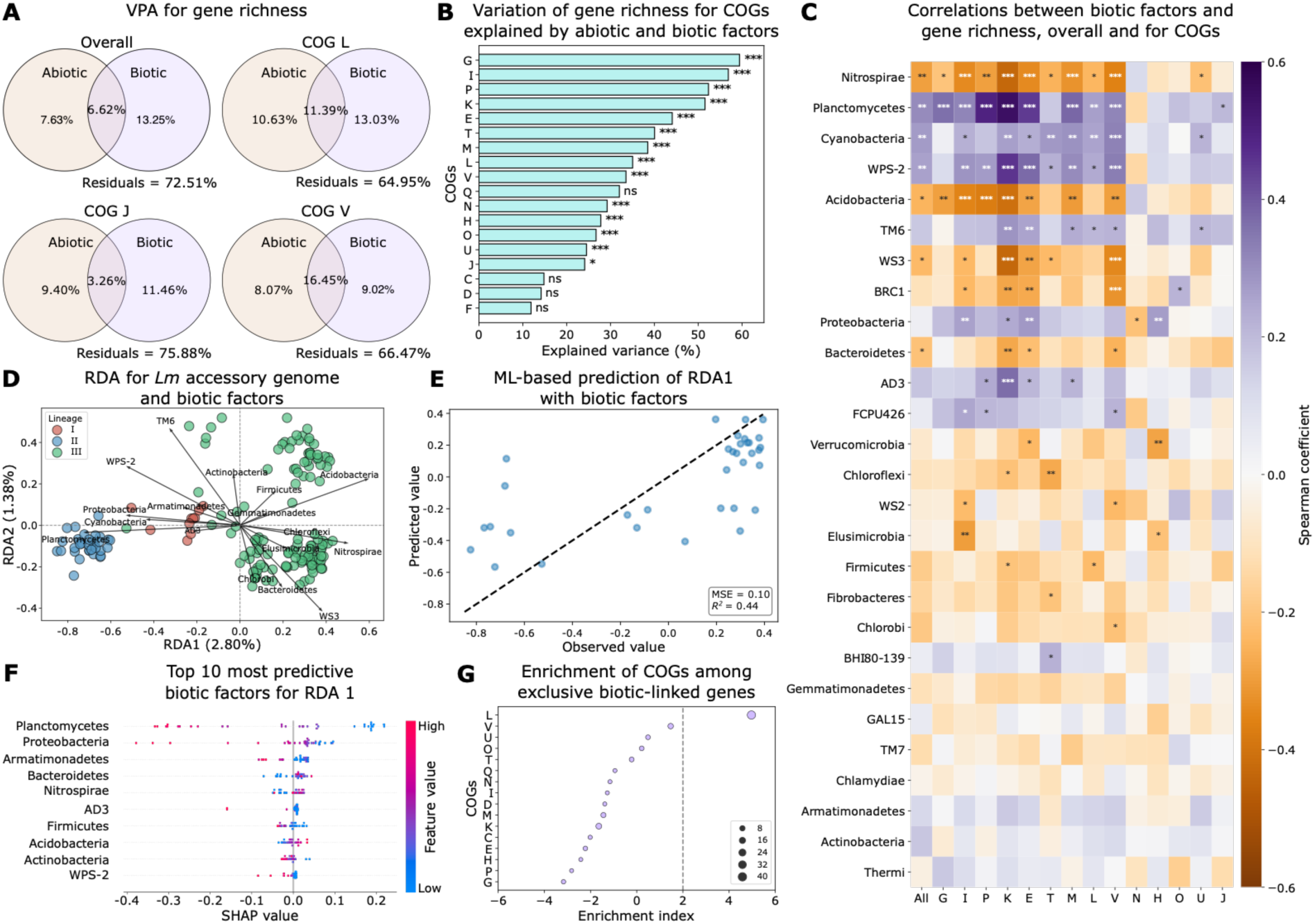
Pangenome fluidity of *Lm* is associated with bacterial community composition. **(A)** Venn diagram of VPA showing the variation of gene richness explained by abiotic and biotic factors. Biotic factors are represented by bacterial community composition at the phylum level. COGs where biotic factors explain more variation than abiotic factors are highlighted. Residuals represent unexplained variation. **(B)** Variation of gene richness for COGs explained by abiotic and biotic factors in VPA. Statistical significance was assessed using a one-sided permutation test. **(C)** Spearman’s correlation between the relative abundance of bacterial phyla and gene richness, overall and for significant COGs identified in VPA. Positive and negative correlations are indicated by purple and orange, respectively. **(D)** RDA biplot illustrating the relationships between the *Lm* accessory genome and 17 phyla selected by Lasso regularization. Points represent individual genomes, color-coded by lineages. The length and direction of the arrows indicate the relative importance and correlation of each phylum with the ordination axes. RDA1 and RDA2 account for 2.80% and 1.38% of the variation of the *Lm* accessory genome explained by selected phyla, respectively. **(E)** Prediction of RDA1 axis values with relative abundance of bacterial phyla using random forests. MSE, mean squared error; *R^2^*, coefficient of determination. The dashed line represents the line of perfect agreement (y = x) where predicted values would exactly match observed values. **(F)** The top ten most predictive biotic factors for RDA1 (SHAP-based; X axis), sorted by descending importance. SHAP values indicate the impact of features on ML model output. **(G)** Enrichment of COGs among exclusive biotic-linked genes. An enrichment index greater than two (indicated by the grey dashed line) signifies significant enrichment (*P* < 0.05). Circle size is proportional to the number of genes annotated to each COG. Abbreviations of COGs are described in Methods. For **(B)** and **(C)**, significance levels are denoted by “*”, “**”, “***”, and “ns” for adjusted *P* < 0.05, < 0.01, < 0.001, and > 0.05 (not significant), respectively.

To pinpoint specific bacterial phyla associated with the gene richness of *Lm*, Spearman’s correlation analysis was performed on the relative abundance of phyla and overall gene richness as well as that for each COG. Results showed that Planctomycetes, Cyanobacteria and WPS-2 were significantly positively correlated with gene richness, while Nitrospirae, Acidobacteria, WS3, and Bacteroidetes showed a significant negative correlation (adjusted Spearman’s *P* < 0.05 for all; **Fig. 2C**). At the functional level, Nitrospirae, Planctomycetes, Cyanobacteria, WPS-2, and Acidobacteria were significantly correlated with gene richness for over half of the 14 significant COGs identified in VPA (adjusted *P* < 0.05 for all; **Fig. 2C; Supplemental Fig. S4**), suggesting a universal effect on the gene content of *Lm* across multiple functions. In contrast, some phyla tend to have a function-specific effect on gene richness. For example, Fibrobacteres and Chlorobi were only significantly correlated with gene richness for COGs T (signal transduction mechanisms) and V (defense mechanisms), respectively (adjusted *P* < 0.05 for both; **Fig. 2C**).

In addition to gene richness, we further examined the relationships between bacterial community composition and the presence/absence patterns of accessory genes. RDA of 17 out of 27 phyla selected by Lasso regularization showed that 2.80% and 1.38% of the variation in RDA1 and RDA2 of the accessory gene content, respectively, was explained by these selected phyla, with TM6, Planctomycetes, WS3, WPS-2, Acidobacteria, Proteobacteria, Nitrospirae, Bacteroidetes, Cyanobacteria, and Chlorobi exerting a strong effect (**Fig. 2D**). Machine learning models further enhanced the prediction, with 44% and 42% of the variation in RDA1 and RDA2 being explained by phyla using random forests and gradient boosting, respectively (**Fig. 2E; Supplemental Fig. S5A**). Based on SHAP analysis [34], TM6, Planctomycetes, WS3, WPS-2, Acidobacteria, Proteobacteria, Nitrospirae, Bacteroidetes, and Cyanobacteria which were found to have a strong effect in the RDA biplot (**Fig. 2D**), were also among the top 10 most predictive phyla for RDA1 and/or RDA2 (**Fig. 2F; Supplemental Fig. S5B**). Analyses on gene richness and accessory genomes altogether suggest that in addition to abiotic factors, bacterial community composition, particularly Nitrospirae, Planctomycetes, Acidobacteria, and Cyanobacteria, play an important role in pangenome fluidity of *Lm*.

To assess the associations between biotic factors and individual accessory genes, we compared the relative abundance of phyla for *Lm* genomes with and without a given accessory gene. A total of 807 accessory genes were found to be significantly associated with at least one phylum, referred to as “biotic-linked genes” (adjusted Mann-Whitney *U P* < 0.05 for all; **Supplemental Table S2**). Functional enrichment analysis showed that these biotic-linked genes were significantly enriched for cell wall/membrane/envelope biogenesis (M) (**Supplemental Fig. S6A**). Comparing these 807 biotic-linked genes to the 803 abiotic-linked genes, 299 genes (27.0%) and 303 genes (27.4%) were exclusively linked to abiotic factors and biotic factors, respectively (**Supplemental Fig. S6B**). Functional enrichment analysis showed that COGs M (cell wall/membrane/envelope biogenesis) and V (defense mechanisms) were significantly enriched among genes exclusively linked to abiotic factors (**Supplemental Fig. S6C**), while L (replication, recombination, and repair) was significantly enriched among genes exclusively linked to biotic factors (**Fig. 2G**), consistent with the VPA result that biotic factors were found to be more important for gene richness involved in this function than abiotic factors (**Fig. 2A**). These results along with gene richness analysis stratified by COGs suggest that bacterial community composition influence the gene content of *Lm* across multiple functions, with pronounced effects on genes involved in cell envelope formation jointly with abiotic factors and uniquely in replication, recombination, and repair.

### Pangenome variation among *Lm* lineages

The 177 *Lm* isolates were classified into three lineages, lineage I (12 isolates), II (29 isolates), and III (126 isolates). Geographically, lineages I (found in the northern and central regions) and II (found along the eastern seaboard and parts of the west) were more widely distributed, whereas lineage III formed a geographic cluster in the central and eastern areas (**Fig. 3A**). Among the 34 abiotic variables, 22 were significantly different among the three lineages (adjusted Kruskal-Wallis *P* < 0.05 for all; **Supplemental Fig. S7**). Of note, lineage III appears to be more capable of surviving in nutrient-limited conditions compared to lineages I and II, as its habitats had significantly lower levels of organic matter, total nitrogen, sodium, and sulfur (adjusted Mann-Whitney *U P* < 0.05 for all; **Supplemental Fig. S7**). In the multidimensional scaling (MDS) analysis based on the 34 abiotic variables, genomes formed significantly distinct clusters by lineages (permutational multivariate analysis of variance, PERMANOVA, *P* = 0.001, pseudo-*F* = 6.703; **Fig. 3B**), which suggests that *Lm* lineages inhabit significantly different abiotic conditions.

**Figure 3.**
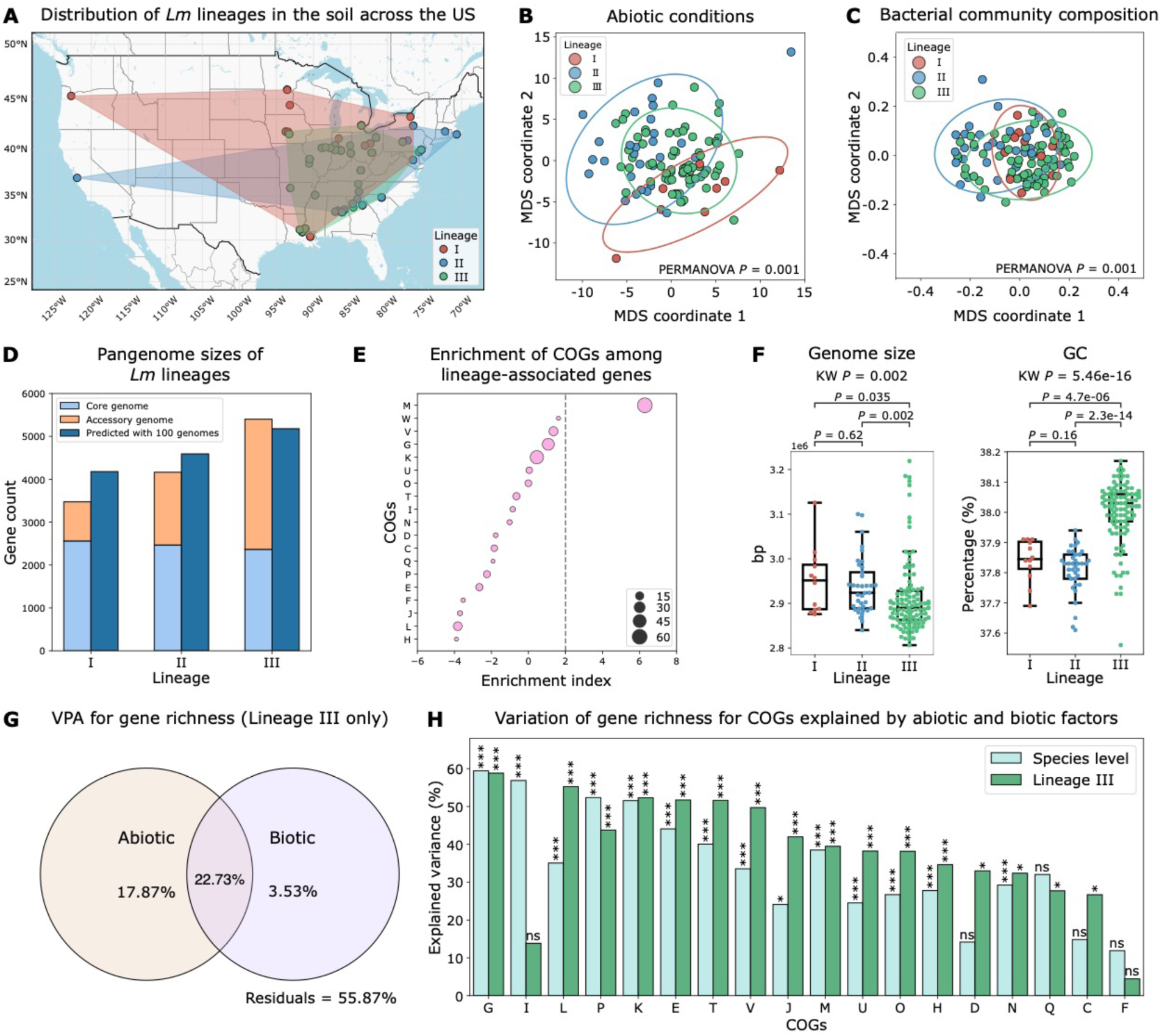
Pangenome variation is substantial among *Lm* lineages. **(A)** Map showing the distribution of *Lm* lineages in soils across the US. Points represent *Lm*-positive samples, color-coded by lineages. Polygons indicate the habitat range of each lineage. This map was adapted from **Extended Data Fig. 5** in Liao et al. 2021 [8]. **(B-C)** Multidimensional scaling (MDS) analysis for *Lm* lineages based on **(B)** abiotic conditions (pseudo-*F* = 6.703) and **(C)** bacterial community composition using weighted UniFrac distances based on OTUs (pseudo-*F* = 8.829). Points represent *Lm* genomes color-coded by lineages. PERMANOVA *P* = 0.001 indicates significant differences in abiotic and biotic conditions among lineages. Ellipse size represents two times the standard deviation from the mean. **(D)** Pangenome sizes of *Lm* lineages, stratified into core and accessory genome sizes, and predicted pangenome sizes based on 100 genomes per lineage. Pangenome size predictions were made using the power law function *cNγ* estimated in **Extended Data Fig. 7** in Liao et al. 2021 [8]. **(E)** Enrichment of COGs among lineage-associated genes. An enrichment index greater than two (indicated by the grey dashed line) signifies significant enrichment (*P* < 0.05). The size of the circles is proportional to the number of genes annotated to each COG. COG abbreviations are described in Methods. **(F)** Genome size and GC content compared among *Lm* lineages. Box plots display the interquartile range (IQR) with the median indicated as a line and whiskers extending to 1.5 times the IQR. Adjusted Kruskal-Wallis (KW) *P* value and adjusted two-sided Mann-Whitney *U P* values are annotated for each boxplot. **(G)** Venn diagram of VPA showing the variation of gene richness for lineage III explained by abiotic and biotic factors. Residuals represent unexplained variation. **(H)** Variation of gene richness for COGs explained by abiotic and biotic factors at the species level (cyan) and within lineage III (green). Significance levels are denoted by “*”, “**”, “***”, and “ns” for adjusted *P* < 0.05, < 0.01, < 0.001, and > 0.05 (not significant), respectively, and “ns” in one-sided permutation tests for VPA.

Similarly, we also identified significant differences in the bacterial community composition compared among *Lm* lineages. Of the 28 phyla, Nitrospirae, Planctomycetes, Acidobacteria, and Proteobacteria showed significantly different relative abundance among the three lineages (adjusted Kruskal-Wallis *P* < 0.05 for all; **Supplemental Fig. S8**). Compared to other lineages, the relative abundance of Planctomycetes and Proteobacteria was significantly higher in lineage II and lower in lineage III, respectively (adjusted Mann-Whitney *U P* < 0.05 for all). Acidobacteria and Nitrospirae were significantly more abundant in lineage III than in lineage II (adjusted *P* = 2.0e-06 and 1.3e-06, respectively; **Supplemental Fig. S8**). MDS analysis showed that genomes formed significantly distinct clusters by lineages based on the beta diversity of bacterial communities represented by weighted UniFrac distances (pseudo-*F* = 5.829; PERMANOVA *P* = 0.001; **Fig. 3C**). Overall, these results suggest that *Lm* lineages occupy significant different abiotic and biotic environmental conditions, which may facilitate adaptive genomic variation in *Lm*.

Indeed, we observed large differences in the gene content among the three *Lm* lineages. While their core genome sizes were similar (2,560, 2,468, and 2,366 core genes for lineage I, II, and III, respectively), their accessory genome sizes were substantially different (913, 1,695, and 3,034 accessory genes for lineage I, II, and III, respectively; **Fig. 3D**). Using the power law function estimated from the pangenome curves for these isolates (**Extended Data Fig. 7** in Liao et al. 2021 [8]), we predicted the pangenome size for each lineage given 100 genomes, which was 4,179, 4,592, and 5,178 genes for lineage I, II, and III, respectively (**Fig. 3D**). Using Fisher’s exact tests, we identified 670 accessory genes significantly associated with lineages (adjusted *P* < 0.05 for all; **Supplemental Table S3**), consistent with the RDA result which identified three distinct clusters by lineages based on accessory genes (**Fig. 1D**; **Fig. 2D**). These lineage-associated genes were significantly enriched for cell wall/membrane/envelope biogenesis (M) based on the functional enrichment analysis (**Fig. 3E**). In addition, we compared the genome size, GC content, virulence factors (*Listeria* pathogenicity island [LIPI]-1, −3, and −4, and internalin [*inl*] genes), SSI 1-2, antibiotic resistance genes (ARGs; *fosX, lin, mprF, norB,* and *sul*), and mobile genetic elements (MGEs), including insertion sequences (IS), transposons, prophages, and plasmids, among lineages. Compared to lineages I and II, lineage III had a significantly smaller genome size (adjusted Mann-Whitney *U P* = 0.035 and 0.0022, respectively, **Fig. 3F**) and higher GC content (adjusted *P* = 4.7e-06 and 2.3e-14, respectively, **Fig. 3F**). For virulence genes, *prfA* (required for the transcription of LIPI-1 [35]) and *hly* (encoding the pore-forming cytolysin Listeriolysin O [35]) in LIPI-1, *inl* genes (encoding cell wall-bound internalins [36]), LIPI-3 genes (encoding the hemolytic and cytotoxic Listeriolysin S [LLS] [37]), and LIPI-4 genes (encoding a cellobiose-family phosphotransferase system [38]) all exhibit significant differences in frequency compared among lineages (adjusted Fisher’s exact *P* < 0.05 for all; **Supplemental Fig. S9A**). Of note, SSI-2, which confers resistance to higher pH [31], was significantly overrepresented in lineage III (adjusted Fisher’s exact *P* = 3.0e-11; **Supplemental Fig. S9A**), consistent with the observation the habitats of lineage III had significantly higher pH compared to lineage II (adjusted Mann-Whitney *U P* = 5.8e-05; **Supplemental Fig. S7**). In addition, proportions of IS elements and transposons significantly differed among lineages, with lineage II having a significantly higher proportion (Kruskal-Wallis *P* = 1.2e-13 and 9.8e-08, respectively; **Supplemental Fig. S9B**). No significant differences were observed for ARGs (adjusted Fisher’s exact test *P* > 0.05 for all; **Supplemental Fig. S9A**), prophages (Kruskal-Wallis *P* = 0.88; **Supplemental Fig. S9B**), and plasmids (Kruskal-Wallis *P* = 0.07; data not shown due to low prevalence).

To compare the associations between pangenomes and abiotic and biotic factors among *Lm* lineages, we intended to perform VPA on gene richness stratified by lineages. However, none of the variation was explained for lineages I and II likely due to the limited sample sizes (*n* = 12 and 29 for lineage I and II, respectively). Therefore, we compared the VPA results for lineage III and the ones at the species level. We found that abiotic and biotic factors jointly explained 44.13% of the variation of gene richness for lineage III (one-sided permutation *P* = 0.01; **Fig. 3G**), which is substantially higher than the 27.49% observed for overall gene richness for *Lm* (**Fig. 2A**). Further, VPA stratified by COGs identified 16 COGs for which variation of gene richness was significantly explained by abiotic and biotic factors. Notably, carbohydrate transport and metabolism (G), replication, recombination, and repair (L), transcription (K), amino acid transport and metabolism (E), and signal transduction mechanisms (T) each exhibited over 50% of the variation explained by these factors (adjusted one-sided permutation *P* < 0.05 for all; **Fig. 3H**). Among the 18 COGs, 13 COGs (72.2%) showed a higher proportion of variation explained in lineage III compared to the species-level result (**Fig. 3H**). These results suggest that the influence of abiotic and biotic factors on the gene content of *Lm* is driven by lineage III and this lineage appears to undergo strong adaption to local nutrient-limited conditions.

### Dispersal dynamics of *Lm* lineages

Homogenizing dispersal across geographic locations can facilitate evolutionary processes that increase genetic similarity among bacterial populations (e.g., homologous recombination) [16–18]. In contrast, dispersal limitation can enhance differentiation in genetic material across populations that emerge from localized environmental adaptation [16,17]. To understand the dispersal patterns of *Lm* lineages, we examined the correlations between genetic similarities, measured by average nucleotide identity (ANI), and geographic distances for each lineage. A distance-decay relationship, in which genetic similarities decline with geographic distances, suggests dispersal limitation [39,40]. Results show that lineage I exhibited a weak distance-decay relation (*R^2^*= 0.01, slope = 1.42e-07; **Fig. 4A**), suggesting frequent homogenizing dispersal across locations. In contrast, lineages II and III showed a strong distance-decay relationship (*R^2^*= 0.51 and 0.33, slope = −2.76e-06 and −2.40e-06, respectively; **Fig. 4A**), indicating strong dispersal limitation. These different dispersal patterns could partially explain the more fluid pangenomes of lineages II and III compared to lineage I. The important role of dispersal in pangenome fluidity is also supported by the strong correlations observed between geolocation and variation in gene content (**Fig. 1A**; **Fig. 1D**; **Fig. 1F**).

**Figure 4.**
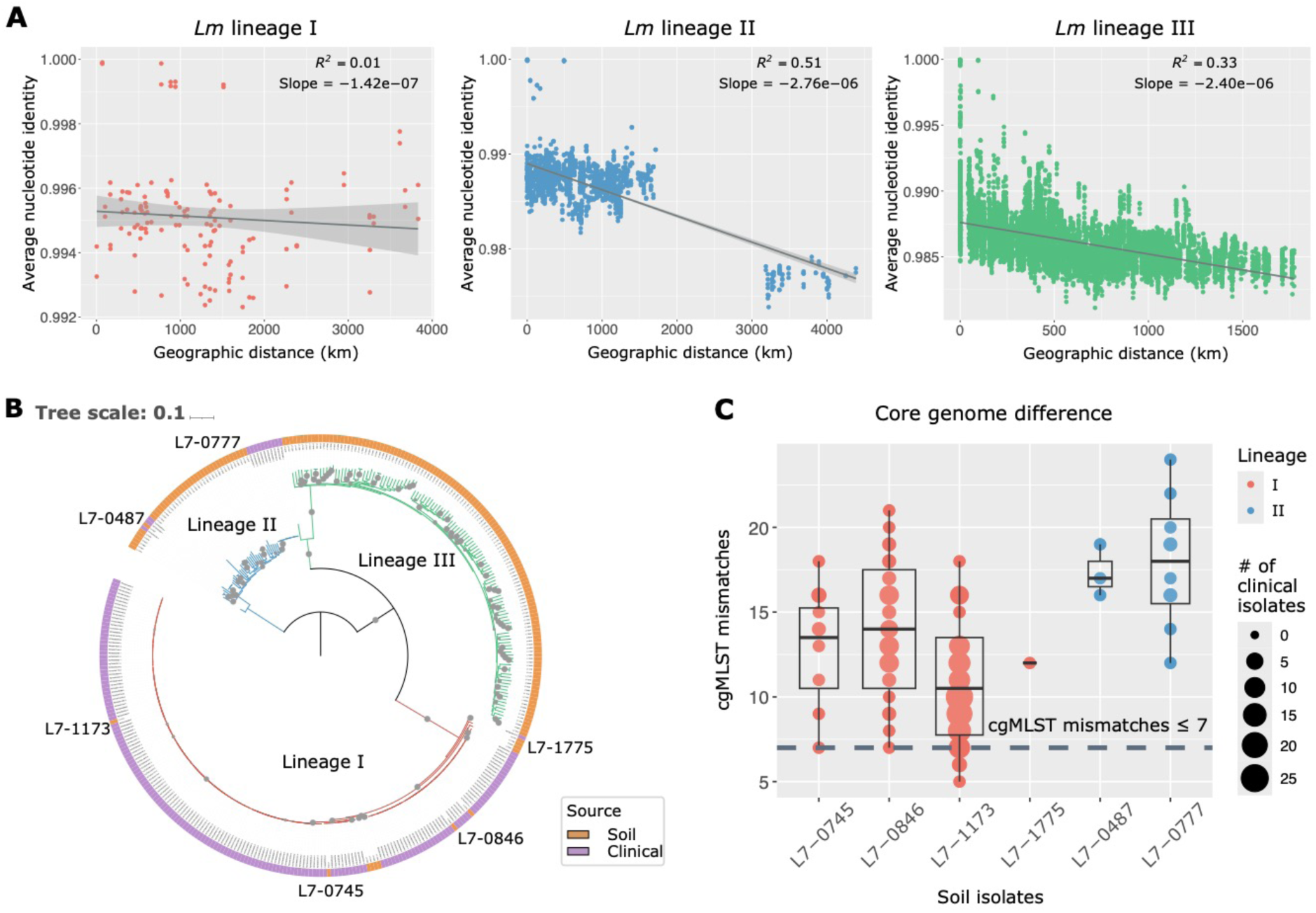
Dispersal patterns vary among *Lm* lineages. **(A)** Linear regression for genetic similarities measured by average nucleotide identity (ANI) and geographic distances for *Lm* lineage I, II, and III. A steeper negative slope of the fitted regression line with a higher *R²* indicates a stronger distance-decay relationship. Shaded areas depict the 95% confidence interval (mean ± 1.96 standard error of mean, SEM) of the linear regression. **(B)** Maximum likelihood phylogenetic tree based on core single nucleotide polymorphisms (SNPs) of 177 soil *Lm* isolates (orange) and 186 closely related clinical isolates (purple), with 1,000 bootstrap replicates. Bootstrap values > 80% are marked by gray circles. The tree is rooted by midpoint, and branches are color-coded by lineages. The six soil isolates with epidemiological links are annotated. **(C)** Core genome multi-locus sequence typing (cgMLST) allelic mismatches between six soil isolates of lineages I (red) and II (blue) and 186 closely related clinical isolates. The horizontal dashed grey line indicates the cutoff (≤ 7 cgMLST mismatches) for documented epidemiological links.[33] The size of the circles is proportional to the number of clinical isolates closely related to a given soil isolate.

Given the frequent homogenizing dispersal potential observed in *Lm* lineage I, we hypothesized that this lineage may have a higher risk of transmission from soils to humans. To test this hypothesis, we compared the genomes of our soil isolates with clinical isolates from human infection downloaded from the National Center for Biotechnology Information (NCBI) Pathogen Detection Isolates Browser [41]. A total of 186 clinical isolates were found to be closely related to six soil isolates (< 50 single-nucleotide polymorphisms [SNPs]) (see Method; **Supplemental Figs. S10 and S11; Supplemental Table S4**). Among these clinical isolates, 173 and 13 were closely related to four soil isolates of lineage I (L7-0745, L7-0846, L7-1173, and L7-1775) and two soil isolates of lineage II (L7-0487 and L7-0777), respectively (**Fig. 4B; Supplemental Figs. S10 and S11; Supplemental Table S4**). Subsequent analysis using core genome multi-locus sequence typing (cgMLST) identified 17 clinical isolates with seven or fewer allelic mismatches with three soil isolates of lineage I (L7-0745, L7-0846, and L7-1173), suggesting epidemiological links (**Fig. 4C**) [33]. Specifically, L7-0745 and L7-0846 each was linked with one clinical isolate, while L7-1173 was linked to 15 clinical isolates (**Supplemental Table S4**). Genomic comparisons, including genome size, GC content, the count of LIPI-1, *inl*, LIPI-3, and LIPI-4 genes, ARGs, and proportions of MGEs (including IS, transposons, and prophages), between these three soil isolates and their corresponding linked clinical isolates further supported the epidemiological links, as no significant differences were identified (Wilcoxon rank-sum *P* > 0.05 for all; **Supplemental Fig. S12**). None of the clinical and soil isolates compared contain any SSI or plasmids. However, it is worth noting that clinical isolates consistently carried more ARG copies than the linked soil isolates (**Supplemental Fig. S12**), suggesting that selective pressures from antibiotics commonly present in the clinical environment enable *Lm* to acquire and retain more ARGs. These findings suggest that *Lm* lineage I has a high risk of transmission from soil to humans.

## DISCUSSION

With in-depth pangenome analysis of soil-dwelling *Lm* that we isolated across the US, this study reveals that the pangenome fluidity of a bacterial pathogen is jointly shaped by environmental selection and dispersal dynamics. We found that selective pressures from abiotic factors acting on *Lm* were mainly imposed by climate (e.g., precipitation and temperature) and soil properties (e.g., aluminum, pH, and molybdenum). Precipitation patterns likely influence bacterial pangenome fluidity by modulating soil moisture, which in turn affects osmotic stress, salinity, and nutrient availability [42,43], potentially selecting for genes that enhance stress tolerance and metabolic flexibility [44]. Likewise, increased temperatures have been reported to impact genetic variation by altering allele frequencies in genes associated with climate adaptation, reducing genetic drift, and promoting homogeneous selection [45,46]. Also, aluminum interacts with soil pH, where under acidic conditions (pH < 5.5), it becomes solubilized into its bioavailable form, which has been found to yield selective pressure on microbial populations [11,47,48]. In addition, in our previous study [8], we identified molybdenum as the second most important environmental factor influencing the nationwide distribution of *Listeria* in soils and observed many of the top genes with SNPs affected by positive selection were involved in the synthesis of folate intermediate molybdopterin, a cofactor that binds molybdenum [49]. Consistent with these findings, we showed that molybdenum is also a critical environmental stressor for gene gain and loss in *Lm* in this study.

While we found that abiotic factors tend to affect gene richness across multiple functions (e.g., carbohydrate, lipid, and inorganic ion transport and metabolism and transcription), the pronounced effects were detected among genes involved in cell envelope synthesis and defense mechanisms. In response to osmotic stress caused by precipitation fluctuations, Gram-positive soil bacteria, such as *B. subtilis, Arthrobacter chlorophenolicus, Rhodococcus erythropolis,* and *Mycobacterium pallens*, were found to modify their membrane composition and structure to maintain cellular homeostasis [50,51]. Also, as the temperature rises, bacterial membranes become more fluid, increasing permeability [52], which can prompt homeoviscous adaptation by adjusting the fatty acid composition of membrane lipids to counteract temperature-induced changes in fluidity [50,52,53]. In addition, under low pH and high aluminum conditions, bacteria can thicken their cell wall by upregulating peptidoglycan synthesis to enhance its tolerance to aluminum toxicity, as observed in *R. erythropolis* [11]. Concurrent enrichment in defense-related genes may help bacteria adapt to other abiotic stresses, such as oxidative stress and desiccation, by encoding stress response proteins crucial for maintaining cellular function and viability under harsh conditions, as observed in *E. coli* [54], *Acidithiobacillus ferrooxidans,* and *Leptospirillum ferriphilum* [55].

In addition to abiotic factors, we found that bacterial community composition also plays a role in the pangenome fluidity of *Lm*. Bacterial phyla that showed the strongest associations were Nitrospirae, Planctomycetes, Acidobacteria, and Cyanobacteria. Nitrospirae, commonly found in oligotrophic environments [56,57], are key players in nitrification, particularly nitrite oxidation, which converts nitrite to nitrate [58]. Nitrate have been found to inhibit the growth of *Lm* [59–61], which may select genetic machinery that involves retaining, modulating, or losing genes in response to the adverse effects. Planctomycetes has been observed co-existing with *Lm* in some environments, such as compost (e.g., soil amendments [62]) and spinach leaves [63], while a negative interaction between *Lm* and Acidobacteria has been reported [23,64,65]. While the understanding of the interactions between Cyanobacteria and *Lm* is limited, it is known that Cyanobacteria could increase the production cyanotoxins (e.g., microcystins and anatoxin) to gain a competitive advantage under nitrogen-/phosphate-imbalance conditions or in the presence of co-occurring microbial species [66–68]. Like abiotic factors, bacterial community composition was associated with genes across multiple functions (e.g., carbohydrate, lipid, and inorganic ion transport and metabolism and transcription). Particularly, the biotic factors appear to impose strong selective pressures on genes involved in cell envelope synthesis jointly with abiotic factors, and uniquely on genes involved in replication, recombination, and repair. Bacterial membranes often house signal transduction systems and histidine kinases crucial for communication, such as the Agr quorum-sensing system and LisRK two-component system [69–71]. These systems regulate the transcription or expression of genes associated with stress response, nutrient uptake, and membrane integrity, enabling bacteria to sense environmental cues and coordinate their behaviors within the community [69–71]. In addition, the association with genes involved in replication, recombination, and repair suggests that potential competitive interactions within the community may expose *Lm* to sources that cause DNA damage [72,73]. In response, *Lm* may initiate DNA repair by activating its SOS system, which is regulated by repair genes such as *recA* [74].

Within-species variation, such as genetic differences among subspecies and phylogenetic lineages, can arise from mutations and gene flow driven by adaptation to abiotic and biotic selective pressures [75]. In *Lm*, substantial pangenome variation was observed among lineages, likely reflecting differences in local environmental adaptation and dispersal dynamics. Lineage III was found to have a highly fluid pangenome but streamlined individual genome size with high GC content, which we attributed to strong adaptation to nutrient-limited conditions (e.g., carbon and nitrogen). Streamlined genomes enable efficient energy use while preserving the flexibility needed to thrive in challenging ecosystems, balancing adaptability with metabolic efficiency [76,77]. Consistent with our observation, a previous study reported that smaller genomes with higher GC content may lower the reproductive cost in carbon-limited soils in bacteria [78]. Environmental stresses have been shown to drive the loss of functionally redundant genes, resulting in genome streamlining, as observed in *Bradyrhizobium diazoefficiens* [79]. In response to varying abiotic and biotic selective pressures encountered in distinct soil conditions across the US, lineage III populations in different locations may lose different sets of genes, increasing the genome variation. In addition to differential local adaptation, strong dispersal limitation observed in this lineage could restrict gene flow across distant populations, enhancing its pangenome fluidity. In contrast, lineage I had a conserved pangenome, likely maintained by frequent homogenizing dispersal. Similar patterns have been observed in other foodborne pathogens, such as *V. parahaemolyticus* and *V. fluvialis*, where effective gene flow facilitated by frequent dispersal contribute to limited divergence and greater population coherence [80,81]. Notably, *Lm* lineage I is commonly associated with produce processing facilities and outbreaks [20,23]. With strong dispersal potential, *Lm* lineage I may spread from soils to nearby produce processing facilities through anthropogenic activities [23,33], eventually infecting humans via direct contact or ingestion, supported by the epidemiological links between soil-derived and clinical isolates observed in this study. These findings underscore the need for lineage-specific source tracking strategies for foodborne outbreaks caused by *Lm*, analogous to the serovar-specific approaches used in *S. enterica* surveillance [82,83].

In summary, we provide critical insights into the ecological mechanisms underlying the bacterial pangenome fluidity and suggest that environmental disturbance, especially shifts in climate patterns, soil properties, and bacterial community composition, can profoundly affect the gene gain and loss process in a bacterial pathogen within its natural reservoir. Specifically, cell surface and defense mechanism-related genes may be targeted by selective pressures from abiotic conditions, while genes involved in replication, recombination, and repair may be uniquely influenced by bacterial communities, in addition to cell surface-related genes jointly linked with abiotic factors. We also highlight the important role of dispersal across geographic locations, along with environmental selection, in shaping within-species pangenome variation. Informed by the dispersal dynamics, we offer critical public health implications, emphasizing the need for targeted genomic monitoring of *Listeria monocytogenes* lineage I in natural environments adjacent to agricultural areas to enhance source tracking of foodborne outbreaks and better manage transmission risks.

## MATERIALS AND METHODS

### Genomic data of soil *Lm* isolates, environmental data, and bacterial community data

Genome assemblies of 177 *Lm* isolates obtained from soil samples collected by us from minimally disturbed natural environments across the US and paired environmental data were analyzed in this study. The genomic and environmental data were published in our previous study investigating the adaptive evolution of *Listeria* [8]. The methods for DNA extraction (using QIAGEN QIAamp DNA MiniKit), whole genome sequencing (via Illumina MiSeq, HiSeq, and NextSeq platforms), and sequencing data processing were detailed in Liao et al. 2021 [8]. Genome assemblies of these isolates were of high quality, with < 300 contigs, an N50 > 50,000, average coverage > 30X, consistent presence of the *sigB* allelic type (AT) in both whole genome and PCR-based assays, no detected contamination using Kraken2 2.0.8 [84], and high genome completeness of 99.99% ± 0.01% (mean ± standard deviation, SD) assessed by CheckM2 [85]. Based on *sigB* allelic types (ATs) and whole genome sequencing data, these *Lm* isolates were classified into lineage I (12 isolates), lineage II (39 isolates), and lineage III (126 isolates). Orthologous genes previously identified using MMseqs2 [86] and annotated for COG functions using eggnog-mapper2 [87] were included in the data analysis in this study. COGs abbreviations are as follows: C: Energy production and conversion; D: Cell cycle control, cell division, chromosome partitioning; E: Amino acid transport and metabolism; F: Nucleotide transport and metabolism; G: Carbohydrate transport and metabolism; H: Coenzyme transport and metabolism; I: Lipid transport and metabolism; J: Translation, ribosomal structure, and biogenesis; K: Transcription; L: Replication, recombination, and repair; M: Cell wall/membrane/envelope biogenesis; N: Cell motility; O: Posttranslational modification, protein turnover, chaperones; P: Inorganic ion transport and metabolism; Q: Secondary metabolites biosynthesis, transport, and catabolism; T: Signal transduction mechanisms; U: Intracellular trafficking, secretion, and vesicular transport; V: Defense mechanisms.

Environmental data analyzed in this study contained 34 variables, including three geolocation variables (longitude, latitude, elevation), 17 soil physicochemical property variables (moisture, total nitrogen [TN], total carbon [TC], pH, organic matter [OM], aluminum [Al], calcium [Ca], copper [Cu], iron [Fe], potassium [K], magnesium [Mg], manganese [Mn], molybdenum [Mo], sodium [Na], phosphorus [P], sulfur [S], and zinc [Zn]), four climate variables (precipitation, wind speed, maximum and minimum temperatures), and 10 land use variables (proportions of open water, barren, forest, shrubland, grassland, cropland, pasture, wetland, and developed [Dev] open space with > 20% and < 20% impervious [IMP] cover in surrounding area). Detailed methods for acquiring these environmental data were detailed in Liao et al. 2021 [8]

Bacterial community data based on 16S rRNA gene amplicon sequencing were published in our previous study comparing the genomic content of *Listeria* populations between soil and food-associated environments [23]. Briefly, total DNA was extracted from soil samples using QIAGEN Power Soil Pro kits, and sequencing was performed on a MiSeq platform with 2 × 250 bp paired-end reads. Raw sequence reads were processed using QIIME2 to remove noises, identify OTUs, classify taxonomy, and calculate beta diversity, including weighted UniFrac distances. Bacterial phyla were referred to as biotic factors in this study.

### Gene richness analysis

VPA was conducted to assess the contributions of **i)** geolocation, soil properties, climate, and surrounding land use, and **ii)** abiotic factors (i.e., all four groups in **i)** combined) and biotic factors (i.e., the relative abundance of bacterial phyla) to the variation of gene richness overall and for COGs at both species and lineage levels. VPA was computed and the adjusted *R^2^* values were visualized in a Venn diagram using the ‘varpart’ function in the vegan package 2.6-4 in R. To assess the significance of the explained variation, a permutation test was performed, in which gene richness was randomly shuffled across genomes 100 times to generate a null distribution for VPA and the resulting explained variation (i.e., expected explained variation) was compared against the observed explained variation using a one-sided test, with significance set at *P* < 0.05. Spearman’s correlation was further conducted to evaluate associations between individual abiotic and biotic factors and gene richness overall and for COGs. To account for multiple testing, *P*-values were adjusted using the Benjamini-Hochberg false discovery rate (BH-FDR) method, with adjusted *P* < 0.05 considered significant.

### RDA, machine learning models, and identification of abiotic/biotic-linked genes

To select between RDA and canonical correspondence analysis (CCA) for exploring relationships between abiotic/biotic variables and accessory genomes, we performed detrended correspondence analysis (DCA) [88]. The first DCA axis length was 0.025, shorter than three standard deviations, indicating that the data is relatively homogeneous and that linear methods, like RDA, are more appropriate for our dataset [88]. Prior to conducting RDA, abiotic factors were standardized to account for the different units of measurement. L1 regularization (Lasso) was applied for feature selection to reduce multicollinearity and enhance model interpretability [89]. The Lasso procedure selected 31 out of 34 abiotic factors and 17 out of 27 bacterial phyla important for the *Lm* accessory genome, which were included in RDA. The length and direction of arrows in the RDA biplot indicate the relative importance and correlation of each abiotic/biotic factor with the ordination axes. DCA and RDA were performed using the ‘decorana’ function in the vegan package 2.6-4 in R and the ‘skbio.stats.ordination.rda’ function in the skbio library in Python 3.6.8, respectively.

To predict the RDA ordination axes from abiotic and biotic factors separately, we implemented a machine learning-based framework using Python 3.6.8 reported in our previous study [90]. In brief, the data were preprocessed and split into training (80%) and testing (20%) sets. The training set was further partitioned into five folds for cross-validation, during which various regression models were trained and evaluated with optimized hyperparameters (see **Supplemental Table S5**). To identify the best-performing model, we conducted a random search across 100 parameter configurations for each algorithm, including regressors based on random forests, neural networks, gradient boosting, support vector machines, and *k*-nearest neighbors. The model with the highest *R^2^* score was then fine-tuned through exhaustive hyperparameter exploration to achieve optimal performance. The importance of the features to the prediction was quantified using SHAP [34].

Two-sided Mann-Whitney *U* tests followed by BH-FDR adjustment were employed to compare abiotic and biotic factors between genomes with and without a given accessory gene. Genes with an adjusted *P* < 0.05 were considered significantly linked to a particular abiotic factor (defined as “abiotic-linked genes”) or linked to a particular phylum (defined as “biotic-linked genes”). Functional enrichment analysis was conducted using a binomial distribution model described in Liao et al. 2023 [23] to identify COGs significantly enriched among abiotic-linked genes, biotic-linked genes, and genes exclusively linked to abiotic or biotic factors.

### Abiotic and biotic environmental conditions of *Lm* lineages and distance-decay relationship

We re-visualized the previously published spatial distribution map of 177 *Lm* genomes (**Extended Data Fig. 5** in [8]) using the Mercator Projection and the Basemap Matplotlib Toolkit 1.2.1 in Python 3.6.8. Kruskal-Wallis tests followed by BH-FDR correction were performed to identify significant differences in abiotic and biotic factors among *Lm* lineages. For factors with an adjusted Kruskal-Wallis *P* < 0.05, pairwise differences between two lineages were further assessed using two-sided Mann-Whitney *U* tests followed by BH-FDR correction. Additionally, MDS analysis was performed using Euclidean distance for abiotic factors, and weighted UniFrac distance for OTUs to compare abiotic conditions and bacterial community compositions among *Lm* lineages followed by a PERMANOVA test to assess clustering significance, with *P* < 0.05 indicating significant clustering by lineages. The distance–decay relationship was inferred by linear regression between geographical distances (km) and biological similarities (indicated by ANI) between genomes for each lineage. A steeper negative slope of the fitted regression line with a higher *R²* indicates a stronger distance-decay relationship.

### Pangenome features of *Lm* lineages

To predict the pangenome size given 100 genomes for each *Lm* lineage, we applied the power law function *cN^γ^*, calculated in **Extended Data Fig. 7** in [8]. The specific functions used were n_pan_ = 2723*N*^0.093^ for lineage I, n_pan_ = 2681*N*^0.122^ for lineage II, and n_pan_ = 2490*N*^0.159^ for lineage III, where *N* is the number of genomes [8]. Fisher’s exact tests followed by BH-FDR correction were performed in Python 3.6.8 to identify genes having significantly different frequencies among lineages, referred to as “lineage-associated genes” (adjusted *P* < 0.05). Functional enrichment analysis was performed using a binomial distribution model described in Liao et al. 2023 [23] to identify COGs significantly enriched among lineage-associated genes.

Additionally, various genomic elements were analyzed, including genome size and GC content (previously reported in Liao et al. 2021 [8]), virulence factors (LIPI-1, -3, -4 and *inl* genes), SSI-1 and 2 (previously reported in Liao et al. 2023 [23]), as well as ARGs and MGEs, including IS, transposons, prophages, and plasmids (previously reported in Goh et al. 2024 [90]). In brief, virulence factors, SSI, and ARG were previously identified through BLASTN searches against reference databases sourced from the BIGSdb-*Lm* platform [33]. Reference virulence factors include LIPI-1 (*prfA, plcA, hly, mpl, actA, plcB*), LIPI-3 (*llsAGHXBYDP*), LIPI-4 (*LM9005581_70009 - LM9005581_70014*), and *inl* genes (*inlAB, C, E, F, G, H, J, K, I, P*). Reference SSI-1 include *lmo0444 - lmo0448* and SSI-2 include *lin0464 - lin0465*. Reference ARGs includes *aacA4, aadC, aadE, aphA, cat_CHL, dfrD, dfrK, ermB, ermG, fexA, fosX, lin, mprF, lnuA, lnuG, mefA, mphB, msrD, norB, penA, qnrB, str, sul, tetM,* and *tetS*. ISEScan (Xie and Tang 2017), TnFinder [92], PHASTEST [93], and PlasmidFinder [94] were previously used for the prediction of IS, transposons, prophages, and plasmids, respectively. While these genomic elements were previously reported, they were not compared among *Lm* lineages. Here, Kruskal-Wallis tests were used to assess the differences in genome size, GC content, and MGEs among lineages, followed by two-sided Mann-Whitney *U* tests with BH-FDR correction to assess the pairwise differences. Fisher’s exact tests with BH-FDR correction were used to compare the frequency of virulence factors, SSI, and ARGs among lineages. An adjusted *P* < 0.05 was considered statistically significant.

### Identification of epidemiological links between *Lm* soil and clinical isolates

The NCBI Pathogen Detection Isolates Browser [41] was used in 2019 to identify human clinical *Lm* isolates found within the same single-linkage SNP cluster as with one or more of the 177 soil *Lm* isolates included in this study. Single-linkage SNP cluster is defined as a cluster of isolates differing by no more than 50 SNPs [95]. A total of 186 closely related clinical isolates were identified, all of which met the genome assembly quality criteria outlined in Liao et al. 2021 [8] (i.e., < 300 contigs, N50 > 50,000), with a high genome completeness of 99.98% ± 0.01% (mean ± SD, **Supplemental Table S6**). A phylogenetic tree including both soil and closely related clinical isolates was constructed using RAxML-8.2.13 [96] based core SNPs identified using kSNP4 [97]. The GTR + G (General Time Reversible + gamma distribution) model selected by jModelTest [96], along with ascertainment bias correction and 1,000 bootstrap repetitions, was applied in the tree construction. The phylogenetic tree was visualized using the Interactive Tree of Life (iTOL) 6 webserver [98].

To identify epidemiologic links between soil and clinical *Lm* isolates, cgMLST was profiled, in which ATs for cgMLST genes were assigned based on the database available on the BIGSdb-*Lm* platform [33]. Isolates differing by up to seven allelic mismatches in cgMLST profiles were considered epidemiologically linked [33]. Genome size, GC content, and total counts for virulence factors, SSI-1 and 2, ARGs, and normalized abundance of MGEs (i.e., IS, transposons, and prophages) were compared between soil isolate L7-1173 and its epidemiologically linked clinical isolates (15 total) using the Wilcoxon rank-sum test, with *P* < 0.05 considered statistically significant. The other two soil isolates (L7-0745 and L7-0846) were epidemiologically linked to one clinical isolate each, so they were not included in the statistical tests.

## DATA AVAILABILITY

The whole genome sequencing data for 177 *Listeria monocytogenes* soil isolates used in this study have been submitted to the NCBI BioProject database under accession numbers PRJNA514286 and PRJNA561882, along with associated environmental data, which were previously published in Liao et al, 2021 [8]. The 16S rRNA gene amplicon sequencing data used in this study have been submitted to the NCBI BioProject database under accession number PRJNA749132 and were published in Liao et al, 2023 [23] NCBI GenBank accession numbers of the genomic assemblies for clinical Lm isolates identified via the NCBI Pathogen Detection Isolates Browser are provided in **Supplemental Table S6**. All data needed to evaluate the conclusions in the paper are present in the paper and/or the Supplementary Materials. Code to replicate all analyses is publicly available at https://github.com/leaph-lab/Lm_pangenome_MS.

## ACKNOWLEDGMENTS

We are grateful to the members of LEAPH (the Liao laboratory) for their enriching discussions.

## AUTHOR CONTRIBUTIONS

Conceptualization, J.L.; Methodology, Y.-X.G., F.H., H.Z., J.L.; Formal Analysis, Y.-X.G., F.H., H.Z., J.L.; Writing – original draft, Y.-X.G.; Writing – review & editing, J.L.; Visualization, Y.-X.G., F.H., H.Z., J.L.; Supervision, J.L.; Funding Acquisition, J.L.

