## Supplementary Figures 1-12 for "Disentangling the impacts of abiotic and biotic environmental factors and dispersal dynamics on the pangenome fluidity of bacterial pathogens"

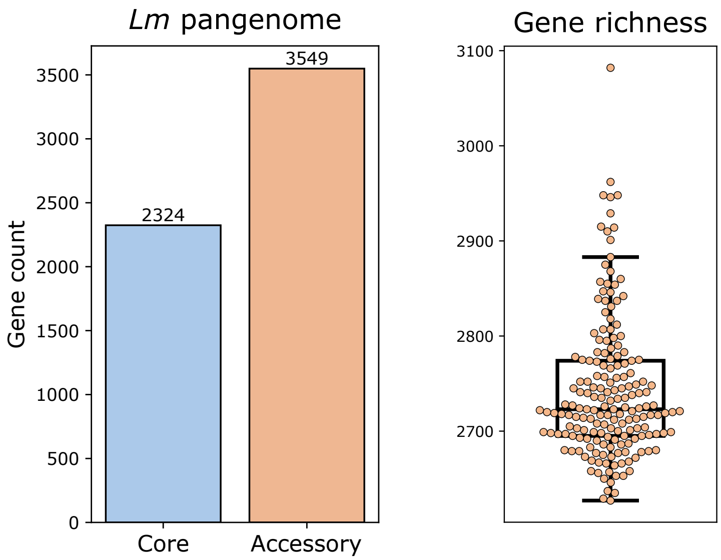


**Supplemental Figure S1. Pangenome size and gene richness of *Lm*.** Pangenome size of *Lm*, stratified into core and accessory genome sizes, and overall gene richness of *Lm*. Box plot displays the interquartile range (IQR) with the median indicated as a line and whiskers extending to 1.5 times the IQR.

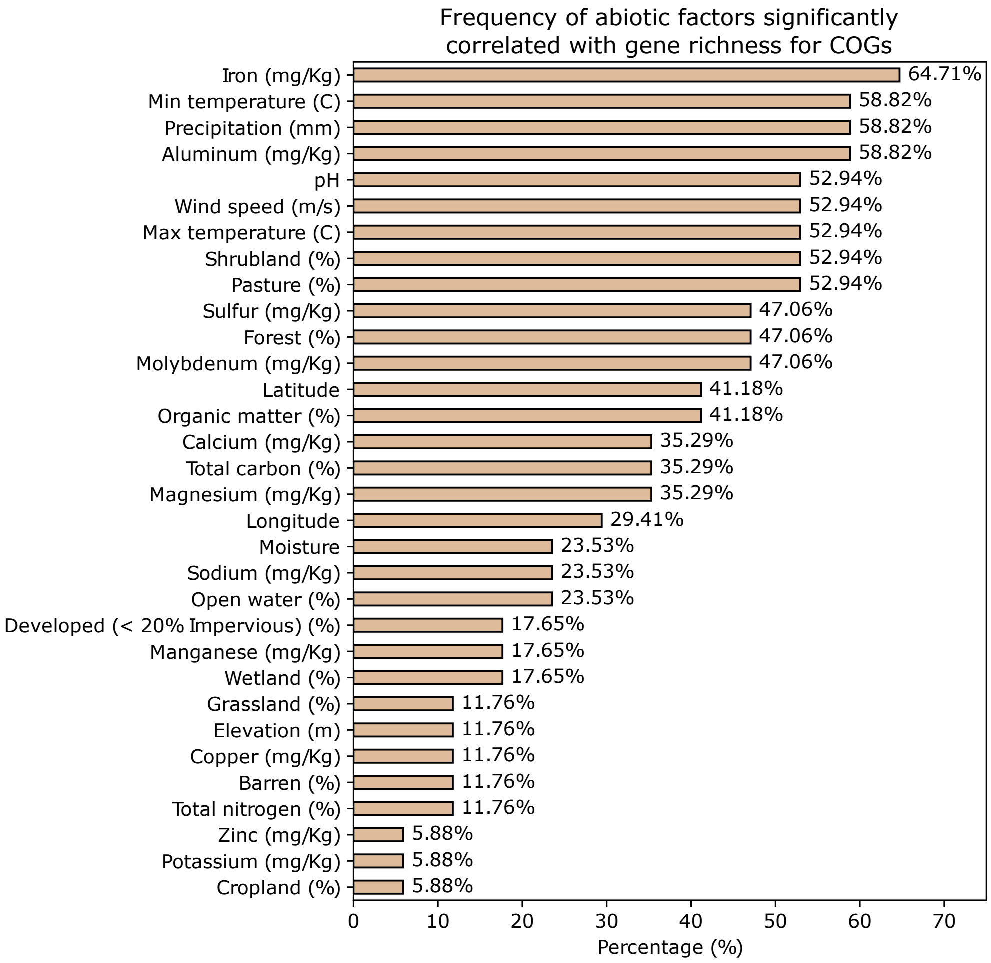


**Supplemental Figure S2. Frequency of abiotic environmental factors significantly correlated with gene richness for Clusters of Orthologous Groups (COGs), sorted in descending order.** Percentage is calculated as the proportion of COGs for which gene richness was significantly correlated with a given abiotic factor among 17 COGs (adjusted Spearman's $P < 0.05$; correlation analysis results are shown in **Fig. 1C**).

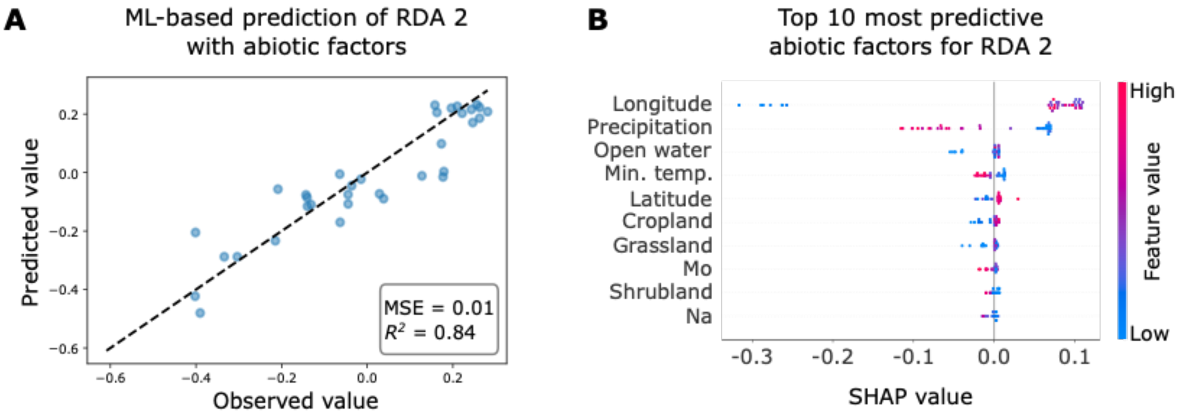


**Supplemental Figure S3. Machine learning (ML) prediction of RDA2 with abiotic factors. (A)** Prediction of RDA2 axis values with abiotic factors using the random forest ML model. MSE, mean squared error; $R^2$, coefficient of determination. The dashed line represents the line of perfect agreement (y = x) where predicted values would exactly match observed values. **(B)** The top ten most predictive abiotic factors for RDA2 (Shapley Additive exPlanations, SHAP-based; X axis), sorted by descending importance. SHAP values indicate the impact of features on ML model output.

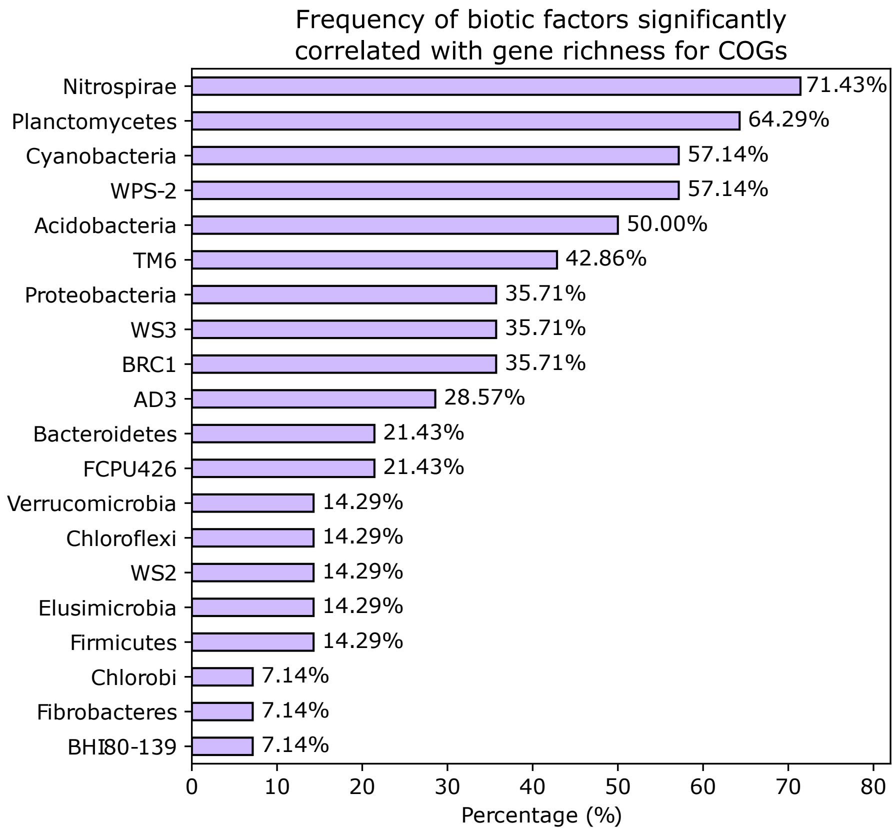


**Supplemental Figure S4. Frequency of biotic factors significantly correlated with gene richness for COGs, sorted in descending order.** Percentage is calculated as the proportion of COGs for which gene richness was significantly correlated with a given bacterial phylum among 14 COGs (adjusted Spearman's *P* < 0.05; correlation analysis results are shown in **Fig. 2C**).

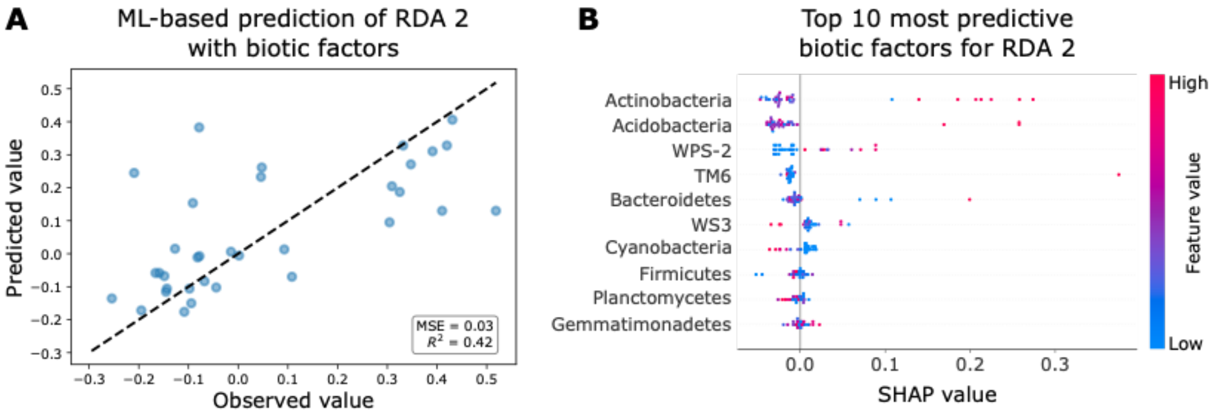


**Supplemental Figure S5. ML-based prediction of RDA2 with biotic factors. (A)** Prediction of RDA2 axis values with relative abundance of bacterial phyla using a gradient boosting model. MSE, mean squared error; $R^2$, coefficient of determination. The dashed line represents the line of perfect agreement (y = x) where predicted values would exactly match observed values. **(B)** The top ten most predictive biotic factors for RDA2 axis values (SHAP-based; X axis), sorted by descending importance. SHAP values indicate the impact of features on ML model output.

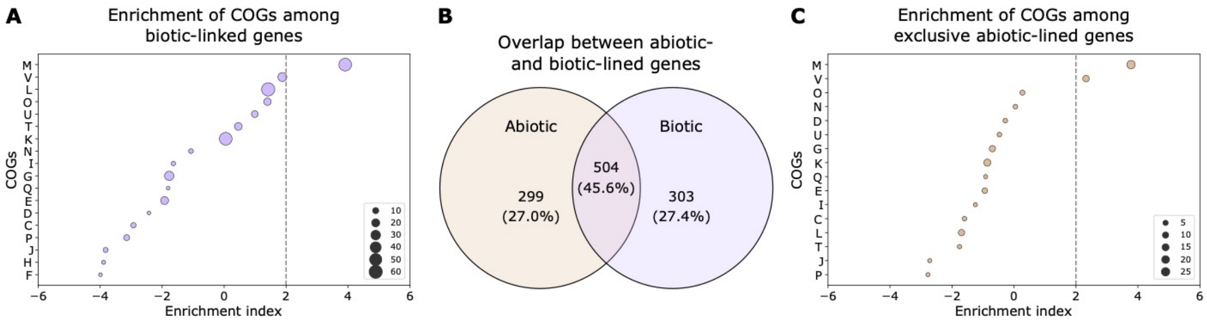


**Supplemental Figure S6. Functions of abiotic- and biotic-linked genes. (A)** Enrichment of COGs among biotic-linked genes. **(B)** Venn diagram showing the overlap between abiotic- and biotic-linked genes. **(C)** Enrichment of COGs among exclusive abiotic-linked genes. For **(A)** and **(C)**, an enrichment index greater than two (indicated by the grey dashed line) signifies significant enrichment (*P* < 0.05). The size of the circles is proportional to the number of genes annotated to each COG. Abbreviations of COGs are described in Methods.

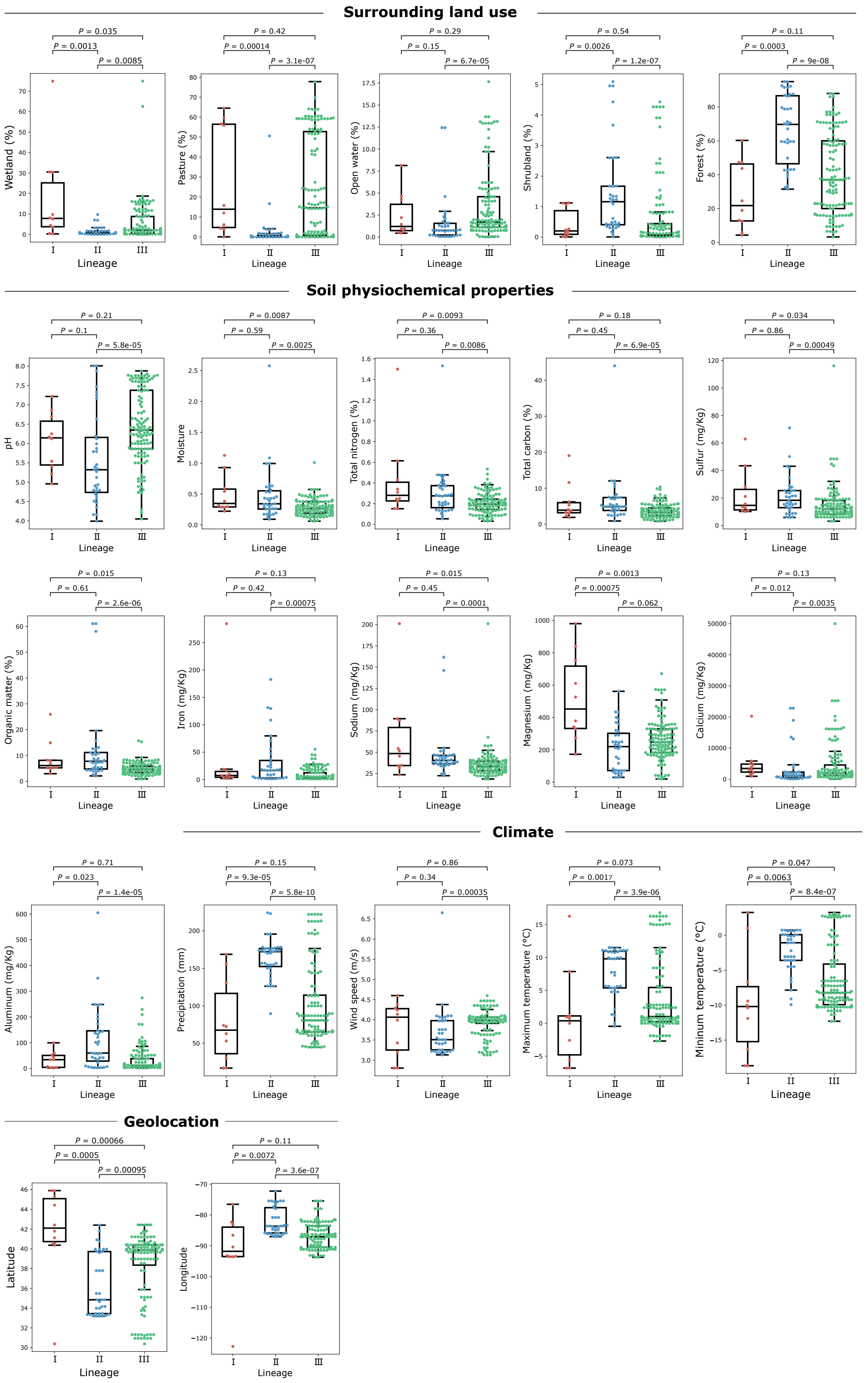
Surrounding land use
P = 0.035
P = 0.0013
P = 0.0085
Wetland (%)
P = 0.42
P = 0.00014
P = 3.1e-07
Pasture (%)
P = 0.29
P = 0.15
P = 6.7e-05
Open water (%)
P = 0.54
P = 0.0026
P = 1.2e-07
Shrubland (%)
P = 0.11
P = 0.0003
P = 9e-08
Forest (%)
Soil physiochemical properties
P = 0.21
P = 0.1
P = 5.8e-05
pH
P = 0.0087
P = 0.59
P = 0.0025
Moisture
P = 0.0093
P = 0.36
P = 0.0086
Total nitrogen (%)
P = 0.18
P = 0.45
P = 6.9e-05
Total carbon (%)
P = 0.034
P = 0.86
P = 0.00049
Sulfur (mg/Kg)
P = 0.015
P = 0.61
P = 2.6e-06
Organic matter (%)
P = 0.13
P = 0.42
P = 0.00075
Iron (mg/Kg)
P = 0.015
P = 0.45
P = 0.0001
Sodium (mg/Kg)
P = 0.0013
P = 0.00075
P = 0.062
Magnesium (mg/Kg)
P = 0.13
P = 0.012
P = 0.0035
Calcium (mg/Kg)
Climate
P = 0.71
P = 0.023
P = 1.4e-05
Aluminum (mg/Kg)
P = 0.15
P = 9.3e-05
P = 5.8e-10
Precipitation (mm)
P = 0.86
P = 0.34
P = 0.00035
Wind speed (m/s)
P = 0.073
P = 0.0017
P = 3.9e-06
Maximum temperature (°C)
P = 0.047
P = 0.0063
P = 8.4e-07
Mininum temperature (°C)
Geolocation
P = 0.00066
P = 0.0005
P = 0.00095
Latitude
P = 0.11
P = 0.0072
P = 3.6e-07
Longitude
Lineage
I
II
III

**Supplemental Figure S7. Abiotic factors compared among *Lm* lineages.** Abiotic factors significantly differing among *Lm* lineages (adjusted Kruskal Wallis $P < 0.05$) are shown. Boxplots display the IQR with the median indicated as a line and whiskers extending to 1.5 times the IQR. Adjusted two-sided Mann-Whitney *U P* values are annotated in each boxplot for pairwise comparison.

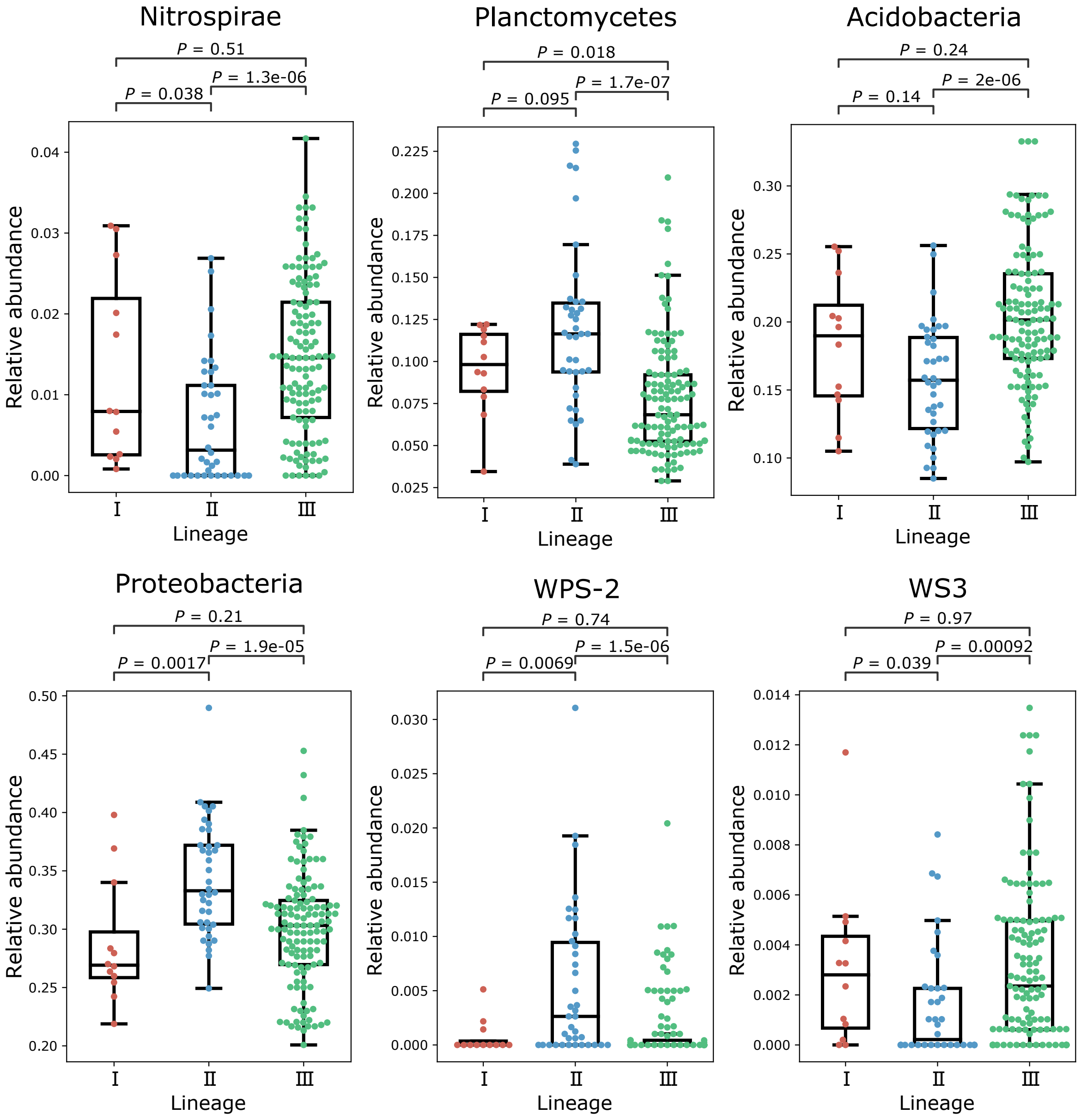


**Supplemental Figure S8. Relative abundance of bacterial phyla compared among *Lm* lineages.** Phyla significantly differing among Lm lineages (adjusted Kruskal Wallis $P < 0.05$) are shown. Boxplots display the IQR with the median indicated as a line and whiskers extending to 1.5 times the IQR. Adjusted two-sided Mann-Whitney *U P* values are annotated in each boxplot for pairwise comparison.

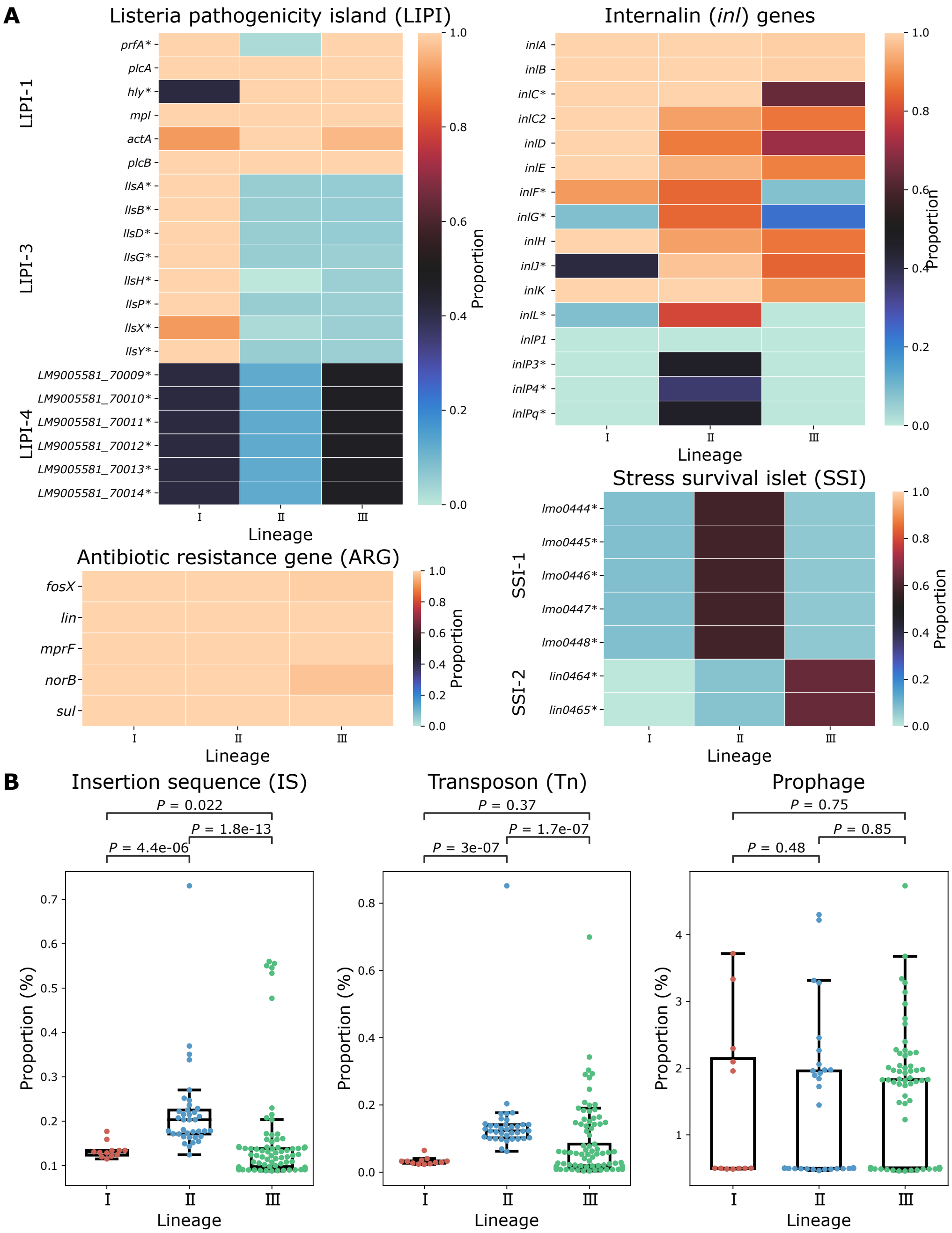


**Supplemental Figure S9. Genetic elements compared among *Lm* lineages. (A)** Prevalence of *Listeria* pathogenicity island (LIPI)-1, -3, and -4 genes, internalin (*inl*) genes, stress survival islet (SSI) 1-2, and antibiotic resistance genes (ARGs) compared among *Lm* lineages. Significance is denoted by '*' for adjusted $P < 0.05$ in a Fisher's exact test. **(B)** Proportion of insertion sequences (IS), transposons, and prophages compared among *Lm* lineages. Boxplots display the IQR with the median indicated as a line and whiskers extending to 1.5 times the IQR. Adjusted two-sided Mann-Whitney *U P* values are annotated in each boxplot for pairwise comparison.

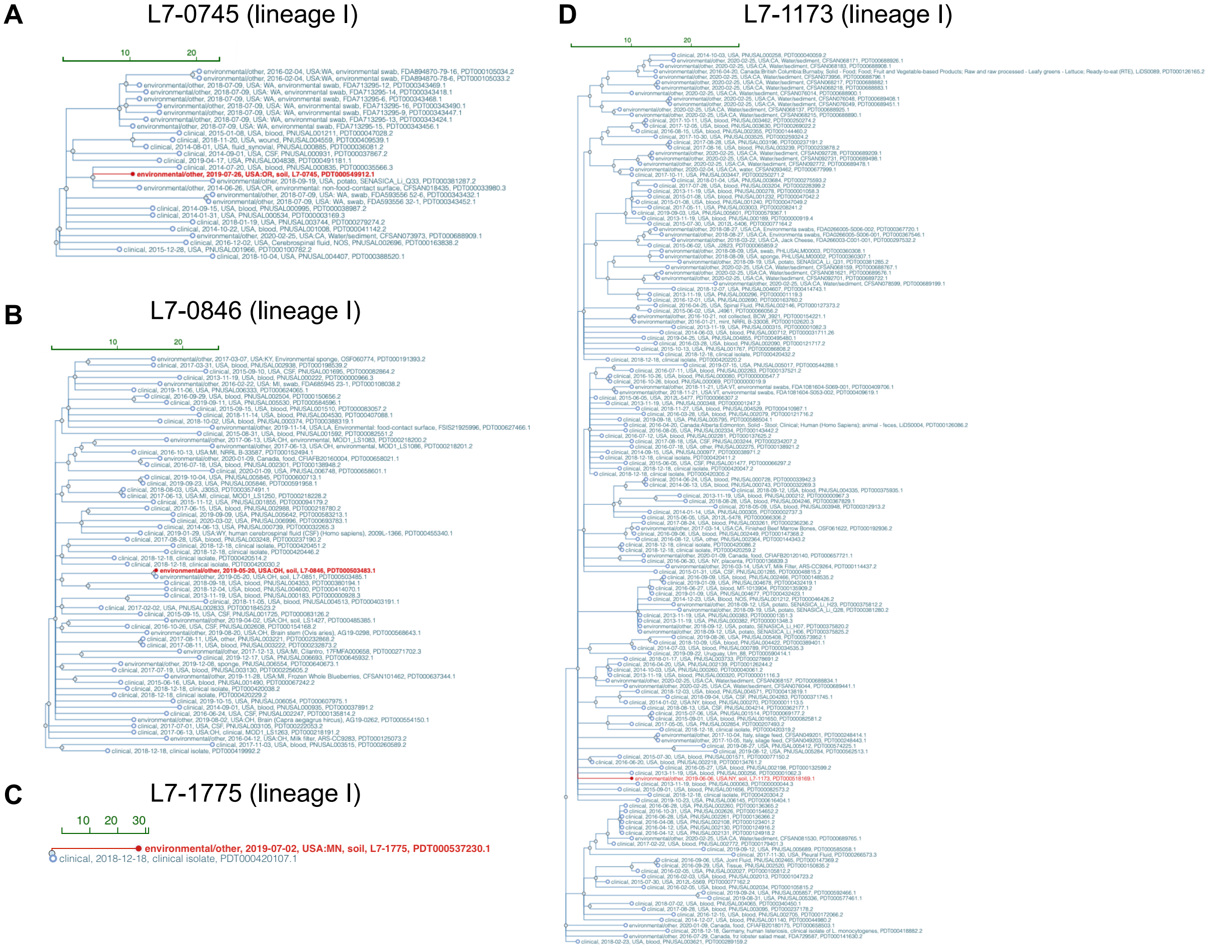


**Supplemental Figure S10. Single-linkage single nucleotide polymorphism (SNP) clusters of closely related soil and clinical *Lm* lineage I isolates. (A-D)** Clinical *Lm* isolates that formed SNP clusters (< 50 SNPs) with *Lm* lineage I soil isolates **(A)** L7-0745, **(B)** L7-0846, **(C)** L7-1775, and **(D)** L7-1173 in this study. Soil isolates are highlighted in red. Trees were downloaded from NCBI Pathogen Detection Isolates Browser.

**A** L7-0487 (lineage II)

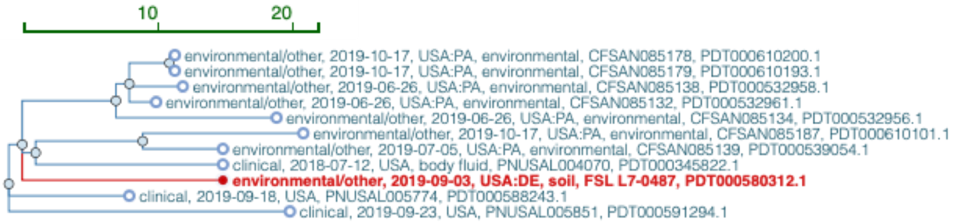


**B** L7-0777 (lineage II)

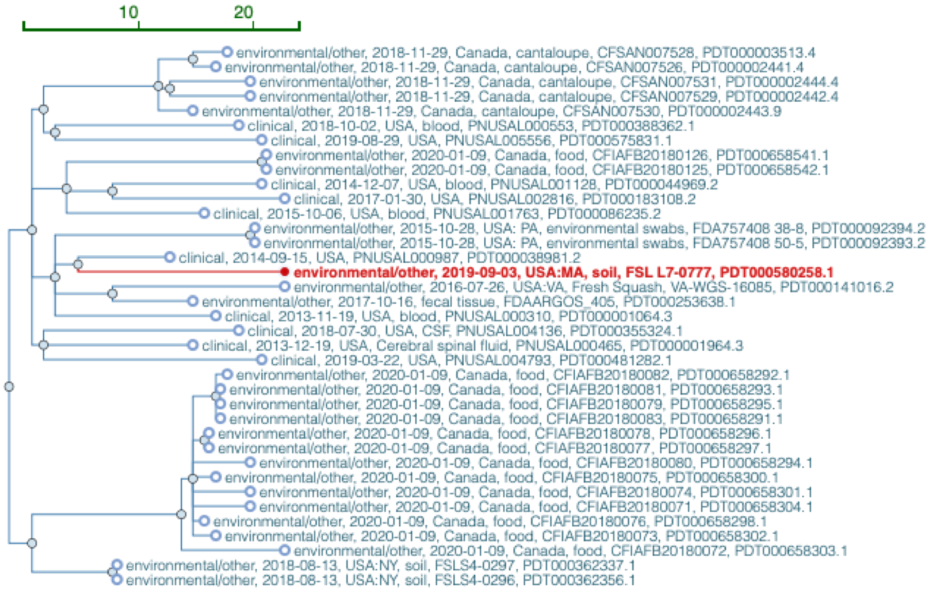


**Supplemental Figure S11. Single-linkage SNP clusters of closely related soil and clinical *Lm* lineage II isolates. (A-B)** Clinical *Lm* isolates that formed SNP clusters (< 50 SNPs) with *Lm* lineage II soil isolates **(A)** L7-0487 and **(B)** L7-0777 in this study. Soil isolates are highlighted in red. Trees were downloaded from NCBI Pathogen Detection Isolates Browser.

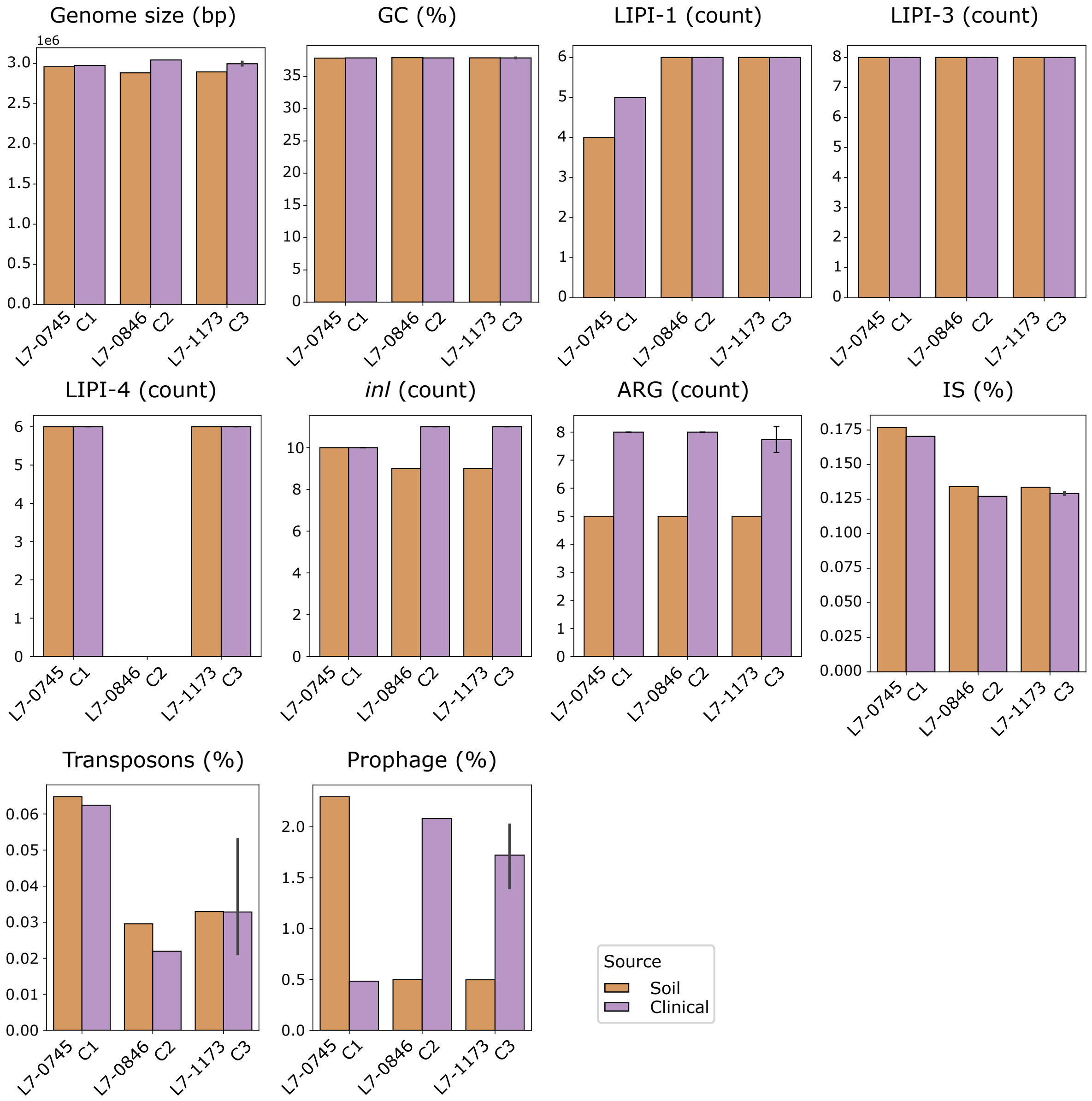


**Supplemental Figure S12. Genomic features compared between epidemiologically linked soil and clinical *Lm* lineage I isolates.** Genomic features, including genome size, GC content, gene counts for LIPI-1, LIPI-3, LIPI-4, *inl*, and ARGs, as well as the proportions of IS, transposons, and prophages, are compared between soil isolates (L7-0745, L7-0846, and L7-1173; orange) and their epidemiologically linked clinical isolates (C1–C3, respectively; purple). Error bars represent the standard deviation.

**Other Supplemental Tables for this manuscript include the following:**

**Supplemental Table S1.** List of abiotic-linked genes, arranged in ascending order of *P*-values.

**Supplemental Table S2.** List of biotic-linked genes, arranged in ascending order of *P*-values.

**Supplemental Table S3.** List of lineage-associated accessory genes and their respective *P*-values identified through Fisher's exact test.

**Supplemental Table S4.** cgMLST differences between soil and clinical *Lm* isolates.

**Supplemental Table S5.** Hyperparameters for machine learning models.

**Supplemental Table S6.** List of clinical *Lm* isolates closely related to soil isolates included in this study, identified via NCBI Pathogen Detection Isolates Browser.
